# DDX41 dissolves G-quadruplexes to maintain erythroid genome integrity and prevent cGAS-mediated cell death

**DOI:** 10.1101/2024.10.14.617891

**Authors:** Honghao Bi, Kehan Ren, Pan Wang, Ermin Li, Xu Han, Wen Wang, Jing Yang, Inci Aydemir, Kara Tao, Lucy Godley, Yan Liu, Vipul Shukla, Elizabeth T. Bartom, Yuefeng Tang, Lionel Blanc, Madina Sukhanova, Peng Ji

**Affiliations:** Department of Pathology, Feinberg School of Medicine, Northwestern University, Chicago, IL; Robert H. Lurie Comprehensive Cancer Center, Northwestern University, Chicago, IL; Galter Health Sciences Library and Learning Center, Feinberg School of Medicine, Northwestern University, Chicago, IL; Division of Hematology and Oncology, Department of Medicine, Feinberg School of Medicine, Northwestern University, Chicago, IL; Department of Cell and Developmental Biology, Feinberg School of Medicine, Northwestern University, Chicago, IL; Center for Human Immunobiology, Feinberg School of Medicine, Northwestern University, Chicago, IL; Department of Biochemistry and Molecular Genetics, Feinberg School of Medicine, Northwestern University, Chicago, IL; Institute of Molecular Medicine, Feinstein Institutes for Medical Research, Manhasset, NY

## Abstract

Deleterious germline *DDX41* variants constitute the most common inherited predisposition disorder linked to myeloid neoplasms (MNs). The role of DDX41 in hematopoiesis and how its germline and somatic mutations contribute to MNs remain unclear. Here we show that DDX41 is essential for erythropoiesis but dispensable for the development of other hematopoietic lineages. Using stage-specific Cre models for erythropoiesis, we reveal that Ddx41 knockout in early erythropoiesis is embryonically lethal, while knockout in late-stage terminal erythropoiesis allows mice to survive with normal blood counts. DDX41 deficiency induces a significant upregulation of G-quadruplexes (G4), noncanonical DNA structures that tend to accumulate in the early stages of erythroid precursors. We show that DDX41 co-localizes with G4 on the erythroid genome. DDX41 directly binds to and dissolves G4, which is significantly compromised in MN-associated *DDX41* mutants. Accumulation of G4 by DDX41 deficiency induces erythroid genome instability, defects in ribosomal biogenesis, and upregulation of p53. However, p53 deficiency does not rescue the embryonic death of Ddx41 hematopoietic-specific knockout mice. In parallel, genome instability also activates the cGas-Sting pathway, which is detrimental to survival since cGas-deficient and hematopoietic-specific Ddx41 knockout mice are viable without detectable hematologic phenotypes, although these mice continue to show erythroid ribosomal defects and upregulation of p53. These findings are further supported by data from a *DDX41* mutated MN patient and human iPSC-derived bone marrow organoids. Our study establishes DDX41 as a G4 dissolver, essential for erythroid genome stability and suppressing the cGAS-STING pathway.

## Introduction

Germline mutations in DDX41 predispose 2-5% of patients with myelodysplastic syndromes (MDS) or acute myeloid leukemia (AML)^1–5^. Patients with *DDX41* mutation tend to be late onset with long-term preceding indolent or mild cytopenia^4,6–8^. More than 80 distinct *DDX41* germline and somatic variants have been reported, making *DDX41* one of the most common MDS/AML predisposition genes^3^. DDX41 is an ATP-dependent DNA/RNA helicase and belongs to the DEAD box family of proteins^9^. It also binds to dsDNA to induce innate immune responses during viral infection^10,11^. Mouse and human DDX41 proteins are highly conserved. DDX41 is functionally important in multiple processes, including mRNA splicing, innate immunity, and rRNA processing^9^. Although clinical evidence shows clear pathologic roles of *DDX41* germline and somatic mutations in myeloid neoplasms, often leading to DDX41 loss of function, the mechanism remains elusive. Previous research on DDX41 led to varied findings. A recent mouse genetic study revealed that biallelic *Ddx41* mutations disrupt snoRNA biogenesis and are incompatible with proliferating hematopoietic cells^12^. Conversely, another study showed that *ddx41* deficiency in zebrafish caused R-loop accumulation, resulting in the expansion of the hematopoietic stem and progenitor cells (HSPCs)^13^. Understanding DDX41’s mechanisms across hematopoietic lineages is crucial for developing innovative therapies for DDX41-mutated MNs.

G-quadruplexes (G4) are four-stranded, noncanonical secondary DNA structures formed in guanine-rich sequences^14^. G4 was reported to be related to the R-loop formation^15,16^. Both structures show accumulation in G-rich regions and cause DNA double-strand breaks (DSBs) by inducing genome instability^13^. DEAD box family proteins, such as DDX5, were shown to be essential to resolve DNA G4^9,17^. G4 homeostasis is also critical for transcriptional regulation during development^18^. Through in vivo studies and human models, we show that DDX41 dissolves G4 structures in the erythroid genome. DDX41 deficiency leads to G4 accumulation, causing genome instability, ribosomal defects, and cGAS-mediated cell death, impairing erythropoiesis and contributing to myeloid neoplasm pathogenesis.

## Results

### Ddx41 is essential for early-stage terminal erythropoiesis

Loss of Ddx41 in mouse hematopoietic cells in vivo was shown to be embryonically lethal with unclear cellular and molecular mechanisms^12^. To investigate Ddx41’s functions in hematopoiesis, we first generated a hematopoietic-specific *Ddx41* knockout mouse model by crossing floxed *Ddx41* (*Ddx41^fl/fl^*) with *VavCre* mice (**Supplemental Figure 1A**). As reported, Ddx41 deficiency led to embryonic lethality^12^. Morphologic examination at E14.5 showed that *VavCre:Ddx41^fl/fl^* embryo was severely pale, indicating defects in erythropoiesis (**Supplemental Figure 1B**). We next purified Ter119 (a mature red cell marker) negative fetal liver HSPCs from the mutant fetuses and cultured the cells in an erythropoietin (Epo)-containing erythroid differentiation system^19,20^. We found that loss of Ddx41 completely blocked erythroid differentiation with a marked increase in cell death (**Supplemental Figure 1C and 1D**), demonstrating that Ddx41 is essential for erythropoiesis.

To study the role of Ddx41 in erythropoiesis in vivo, we generated two erythroid-specific *Ddx41* knockout mouse models, *EpoRCre:Ddx41^fl/fl^*and *HBBCre:Ddx41^fl/fl^* mice, to test the functions of Ddx41 at different stages of erythropoiesis. *EpoRCre:Ddx41^fl/fl^* mice manifest *Ddx41* deletion at the progenitor stages of erythropoiesis (approximately the CFU-E (colony forming unit-erythroid) stage), whereas *HBBCre:Ddx41^fl/fl^* mice exhibit *Ddx41* deletion at the terminal stages of erythroid differentiation^21^ (**Figure 1A and Supplemental Figure 1E**). We found that *EpoRCre:Ddx41^fl/fl^* mice were also embryonically lethal (**Figure 1B**). The lack of appropriate morphogenesis suggests that Ddx41 could be critical for primitive erythropoiesis (**Figure 1C**). We next analyzed *EpoRCre:Ddx41^fl/+^* mice that survived to adulthood. These mice exhibited normal fetal liver erythropoiesis (**Supplemental Figure 2A**). They showed mild macrocytic anemia with normal white blood cell and platelet counts at 2 months old (**Figure 1D and Supplemental Figure 2B)**, which closely mimics patients with *DDX41* germline mutations before the development of overt MNs upon somatic second hit^22^. We sacrificed the *EpoRCre:Ddx41^fl/+^*mice and analyzed bone marrow erythroid cells using flow cytometry of cell surface CD71 and Ter119, two established markers for erythroid maturation. We found significantly compromised terminal erythropoiesis and increased myeloid cells percentage-wise and in absolute numbers with *Ddx41* heterozygosity (**Figure 1E and Supplemental Figure 2C**). This was accompanied by increased Ter119 positive erythroid cells in the spleen (**Figure 1F and Supplemental Figure 2D**), which indicates compensatory extramedullary stress erythropoiesis.

**Figure 1.**
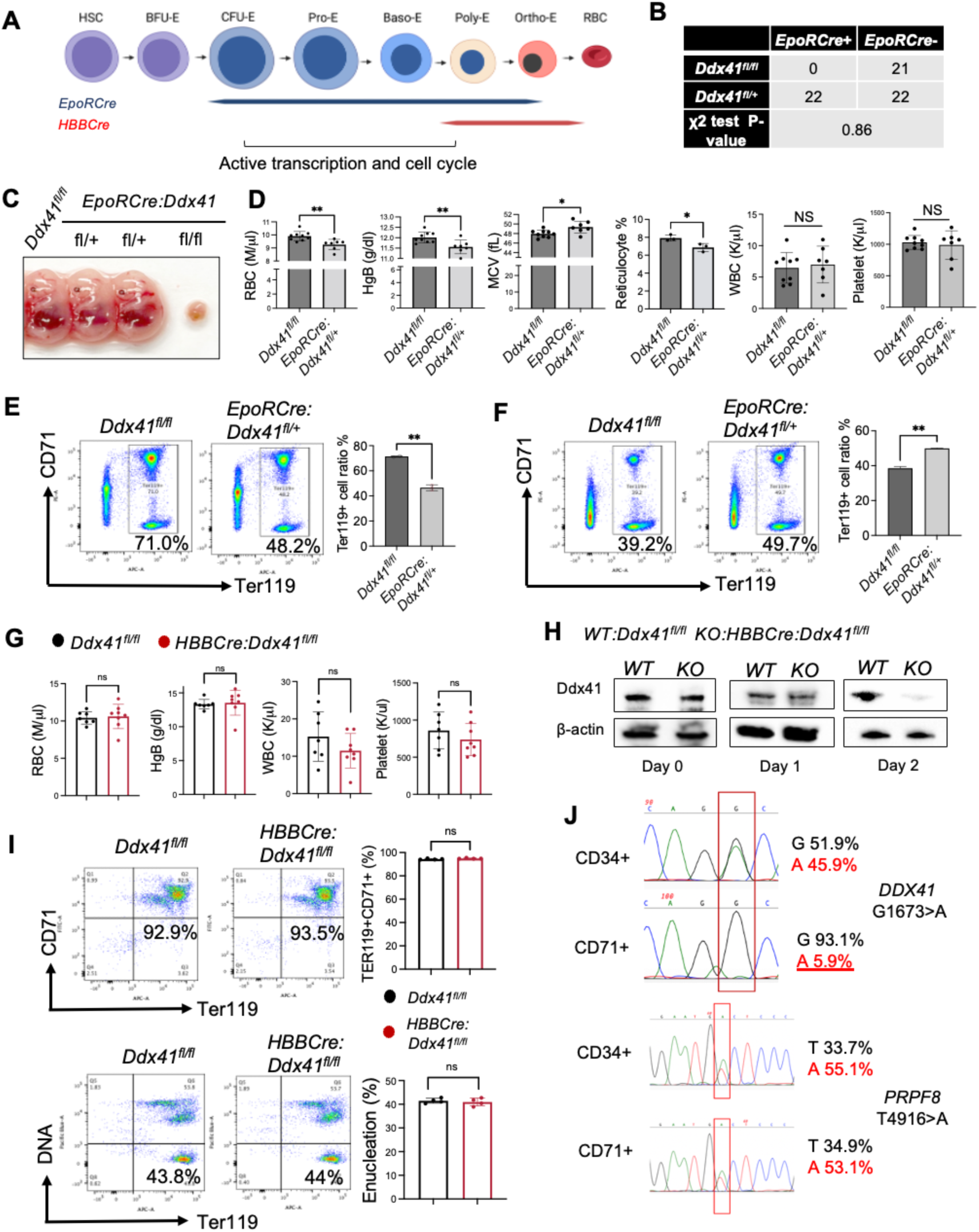
Ddx41 is essential for early-stage terminal erythropoiesis. (A) Schematic illustration of erythropoiesis. Terminal erythropoiesis starts from the stages of CFU-E to proerythroblast (ProE). The stages of EpoR and HBB-driven Cre expression are shown. (B) Mendelian segregation of indicated mice. A χ^2^ test yields a p-value of 0.86, indicating no significant deviation from the expected Mendelian segregation. (C) Representative pictures of E14.5 embryos with the indicated genotypes. (D) Complete blood count (CBC) of the indicated mice at 2 months of age. (E) Flow cytometric assays of bone marrow erythroid cells from the indicated mice from D. Statistical analysis is on the right. (F) Same as E except spleens were analyzed. (G) CBC analysis of indicated mice at 2 months of age. No statistically significant differences were found. (H) Western blotting of Ddx41 from the ex vivo Epo medium-cultured bone marrow erythroid cells from the indicated mice at 2 months of age. (I) Flow cytometric assays of bone marrow erythroid differentiation (upper panels) and enucleation (bottom panels) from the indicated mice in G. No statistically significant differences were found (right). (J) Sanger sequencing of *DDX41* and *PRPF8* in CD34+ HSPCs and CD71+ erythroid progenitor cells purified from bone marrow aspirate of a patient with MDS. *DDX41* mutation is somatic with unclear zygosity. All the error bars represent the SEM of the mean. The comparison between two groups was evaluated with 2 tailed t test. * p<0.05, **p<0.01. ns: not significant.

In contrast, *HBBCre:Ddx41^fl/fl^* mice were viable with no detectable hematologic phenotypes, manifested by normal complete blood count (CBC) at 2 months old (**Figure 1G and Supplemental Figure 2E**). To confirm that *Ddx41* is knocked out in the late-stage terminal erythroblasts in *HBBCre:Ddx41^fl/fl^* mice, we cultured *HBBCre:Ddx41^fl/fl^*bone marrow lineage negative cells in the Epo-containing medium in which mouse erythroid progenitor cells can rapidly differentiate and proliferate to mature red blood cells in 2 days (**Supplemental Figure 2F**). Western blotting demonstrated that Ddx41 showed a slight decrease on day 1 when the cells were predominantly at the early stages of terminal erythropoiesis proerythroblast stages^23–25^. Ddx41 was markedly decreased on day 2 when the cells were at the late-stage terminal erythropoiesis (**Figure 1H**). Consistent with the CBC data, flow cytometry analyses showed no difference in the differentiation and enucleation of the cultured bone marrow erythroid cells from *HBBCre:Ddx41^fl/fl^* and control mice (**Figure 1I**). Similarly, we found no defects in erythropoiesis or the differentiation of other lineages in vivo in the bone marrow and spleen of *HBBCre:Ddx41^fl/fl^* mice (**Supplemental Figure 2G-2K**). Together, these results demonstrate that Ddx41 is critical at the early stages of erythropoiesis but dispensable at the late stages in mice.

We generated additional lineage-specific *Ddx41* knockout mouse lines to study its roles in hematopoiesis. These include *CD11c-Cre:Ddx41^fl/fl^, LysMCre:Ddx41^fl/fl,^ and MRP8Cre:Ddx41^fl/fl^,* which knockout *Ddx41* predominantly in dendritic cells, monocytes, and myeloid/granulocytes, respectively (**Supplemental Table 1**). Interestingly, these mice were all viable with no obvious abnormalities in their CBC at young ages. Flow cytometry studies revealed no obvious differences in the hematopoietic tissues in these animals (**Supplemental Figure 3-5**), demonstrating that Ddx41 is dispensable for the development of these lineages.

Given the significant role of Ddx41 in erythropoiesis in animal models, we wonder whether the same also occurs in MN patients with *DDX41* mutations. We identified an MDS patient carrying a somatic *DDX41* G1673>A mutation (G530D) on the helicase domain, which was indicated to potentially disrupt the ATP binding to DDX41^5,7^. In addition to the *DDX41* mutation, this patient also carries a *PRPF8* T4916>A mutation. The patient has a long-standing MDS with anemia, neutropenia, and occasional mild thrombocytopenia. We purified the patient’s bone marrow CD34+ HSPCs and CD71+ erythroid progenitor cells. Sanger sequencing showed that *DDX41* mutation exists in CD34+ cells but at a noise background level in CD71+ cells. In contrast, the *PRPF8* mutation is present at an equal allele frequency in both CD34+ and CD71+ cells (**Figure 1J**). These findings suggest that the *DDX41* mutation is incompatible with erythroid survival or CD34+ cells fail to differentiate to the erythroid lineage.

### DDX41 controls the level of G-quadruplexes in early erythropoiesis

DDX41’s role in erythropoiesis is underscored by studies in zebrafish, where its deficiency leads to anemia through ineffective erythropoiesis, involving DNA damage response pathways^26^. DDX41 is also critical in regulating DNA secondary structures such as R-loops^13,27^, which frequently co-exist with G-quadruplexes (G4). G4 is known to stabilize R-Loop and regulate transcription by interacting with various chromatin-binding proteins, including several DDX family proteins^14,18,28^. The role of G4 in erythropoiesis is unknown.

To study the role of G4 in erythropoiesis and whether the loss of DDX41 affects G4 formation, we first determined G4 levels in the bone marrow erythroid cells in vivo. We found that G4 is significantly increased in Ter119 positive erythroid cells compared to Ter119 negative cells (**Figure 2A**). We next cultured mouse bone marrow lineage-negative cells in the Epo-containing medium and tested G4 levels at different differentiation stages. We found that G4 significantly increased on day 1, corresponding to the highest proliferation and replication stage (**Figure 2B**). The level of G4 decreased on day 2 when over 30% of the cells were enucleated. The same trend of changes in G4 levels was observed in cultured human CD34+ cells toward erythroid differentiation (**Figure 2C, Supplemental Figure 2F**). Importantly, G4 levels in vivo in the erythroid cells are particularly higher than those in the other hematopoietic lineages (**Figures 2D and 2E**), further indicating a critical role of G4 during erythropoiesis.

**Figure 2.**
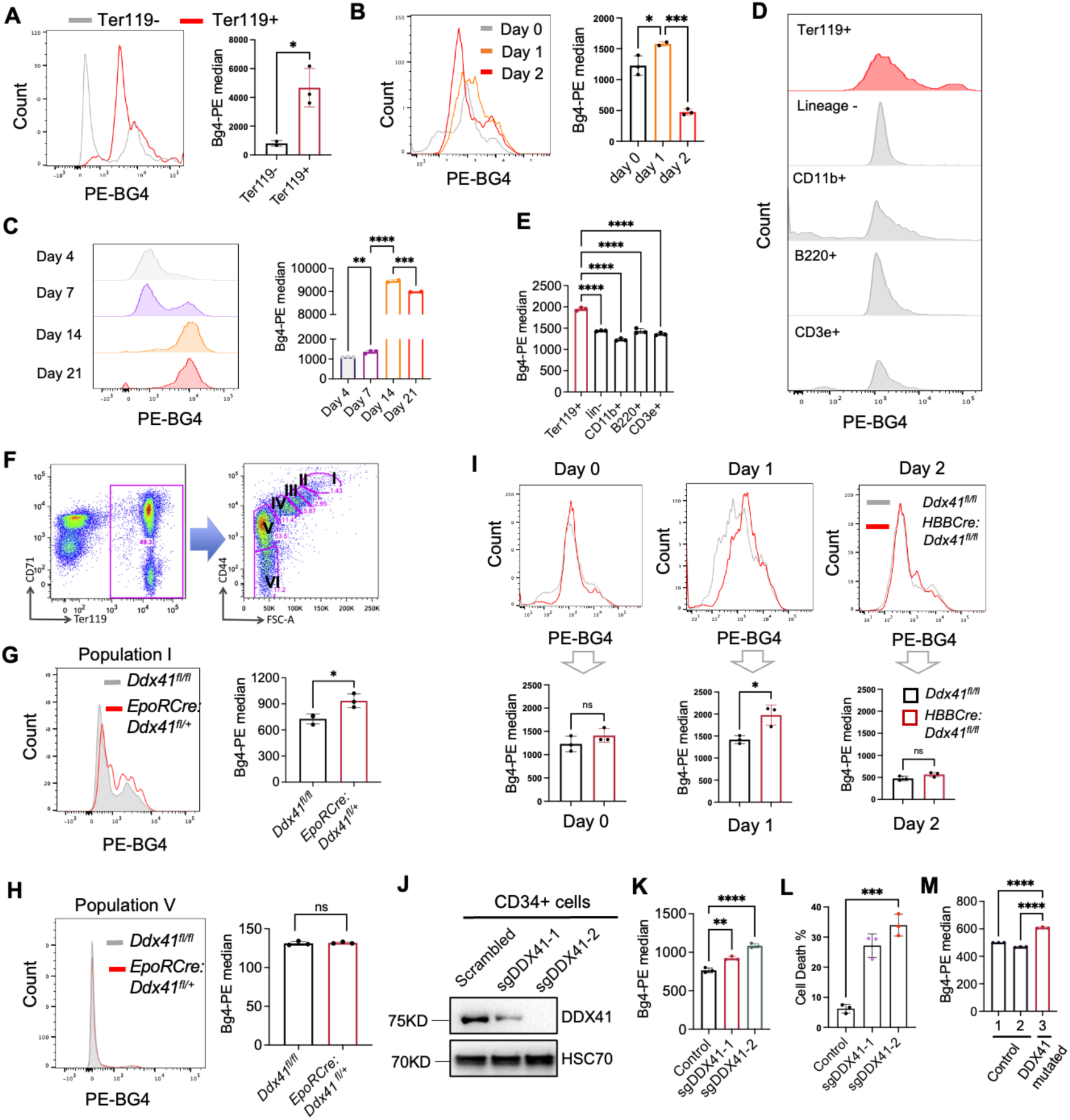
Loss of DDX41 induces accumulation of G-quadruplexes. (A) Ter119 negative cells and Ter119 positive erythroid cells were purified from wild-type mouse bone marrow cells. G4 levels were tested by flow cytometry using the BG4 antibody that specifically recognizes G4. Quantification is on the right. (B) Bone marrow lineage-negative cells were cultured in Epo medium for 2 days. G4 levels were tested on different days using flow cytometry by the BG4 antibody. Quantification is on the right. (C) CD34+ human HSPCs were cultured in Epo medium for 21 days. The levels of G4 were measured by flow cytometry as in B at the indicated time. Cells at day 7, 14, and 21 represent proerythroblasts, polychromatic to orthochromatic erythroblasts, and orthochromatic to mature red blood cells, respectively. (D) Flow cytometric assays of G4 levels in the indicated bone marrow lineage cells purified from wild-type mice. (E) Quantification of D. (F) Gating strategy of various erythroblasts. Populations I to VI represent proerythroblasts, basophilic erythroblasts, polychromatic erythroblasts, orthochromatic erythroblasts, late orthochromatic to reticulocytes, and mature red blood cells, respectively. (G-H) Flow cytometric assay of G4 level in bone marrow erythroid populations I (G) and V (H) from the indicated mice. Quantification is on the right. (I) Bone marrow lineage negative cells from the indicated mice were cultured in Epo medium for 2 days. G4 levels on different days were measured by flow cytometry using BG4 antibody. Quantification is below the histogram. (J) CD34+ cells were transduced with lentiviral vectors expressing indicated sgRNAs and Cas9. Cells were then harvested for Western blotting of the indicated proteins at day 9 in culture. (K) Quantitative analyses of G4 levels in cells from J using flow cytometric assays. (L) Quantitative analyses of cell death in cells from J using flow cytometric assays. The dead cells are defined as propidium iodide and annexin V double positive. (M) Quantitative analyses of G4 levels in bone marrow mononuclear cells from the patient with DDX41 mutated MDS. All the error bars represent the SEM of the mean. The comparison between two groups was evaluated with 2 tailed t tests, and the comparison among multiple groups was evaluated with 1-way ANOVA tests. * p<0.05, **p<0.01, ***p<0.001, and ****p<0.0001. ns: not significant.

Since *VavCre:Ddx41^fl/fl^* mice are embryonically lethal, we used *EpoRCre:Ddx41^fl/+^* and *HBBCre:Ddx41^fl/fl^*mice to investigate Ddx41’s role in G4 accumulation in vivo. We first analyzed G4 levels in the bone marrow erythroid cells in *EpoRCre:Ddx41^fl/+^* mice using a Ter119 and CD44-based gating strategy in which different stages of erythroid precursors can be separated based on CD44 expression (**Figure 2F**)^29,30^. We found a mild but statistically significant increase in G4 level in the proerythroblast population (population I) in *EpoRCre:Ddx41^fl/+^* mice compared to their wild-type counterparts (**Figure 2G**). G4 levels in the late-stage orthochromatic erythroblasts (population V) showed no difference (**Figure 2H**). In *HBBCre:Ddx41^fl/fl^*mice, we did not find significant differences among different populations in the same assay (data not shown), which is consistent with the lack of phenotypes in these mice. It is also difficult to pinpoint what developmental stage(s) G4 starts accumulating in vivo in these mice since most of the erythroid cells in the marrow are at the late stages of terminal erythropoiesis^31^ when G4 level is reduced. Therefore, we purified lineage-negative HSPCs from these mice and cultured them in vitro in Epo-medium. As Ddx41 started to reduce on day 1 (**Figure 1H**), the G4 level started to increase. The G4 level reduced back to the normal range on day 2, as expected (**Figure 2I**). These results are consistent with the phenotypes of these mice and the adverse effect of increased G4 in early, but not late, stages of erythropoiesis. To further demonstrate the role of DDX41 in reducing G4 levels in human erythroid cells, we knocked out DDX41 through CRSPR/Cas9 in CD34+ human HSPCs using two different sgRNAs. Indeed, this led to a significant reduction in DDX41 protein levels (**Figure 2J**) and a marked increase in G4 levels and cell death (**Figure 2K and 2L**). Consistent with these model systems, we also observed increased G4 levels in the bone marrow mononuclear cells of the MN patient with DDX41 mutation (**Figure 2M**).

We next applied pyridostatin (PDS), a selective G4 binding small molecule that stabilizes G4 and perturbs G4 homeostasis^32^, in our in vitro erythroid differentiation system. PDS significantly increased G4 levels in cultured day 1 erythroid cells (**Figure 3A**), induced dosage-dependent inhibition of erythroid differentiation, and increased cell death (**Figure 3B**). Treatment of wild-type mice with a three-day high dose of PDS also significantly compromised bone marrow erythropoiesis in vivo (**Figure 3C**), although no anemia was observed due to the long half-life of red blood cells. We next treated wild-type mice with PDS for 6 weeks through subcutaneously implanted pumps that chronically release the compound. Indeed, this treatment led to significant anemia (**Figure 3D**). The white blood cell count was also reduced, primarily due to PDS-induced lymphopenia (**Figures 3D and 3E**). As expected, chronic PDS treatment also led to ineffective terminal erythropoiesis and proportionally increased myeloid cells in the bone marrow (**Figures 3F and 3G**).

**Figure 3.**
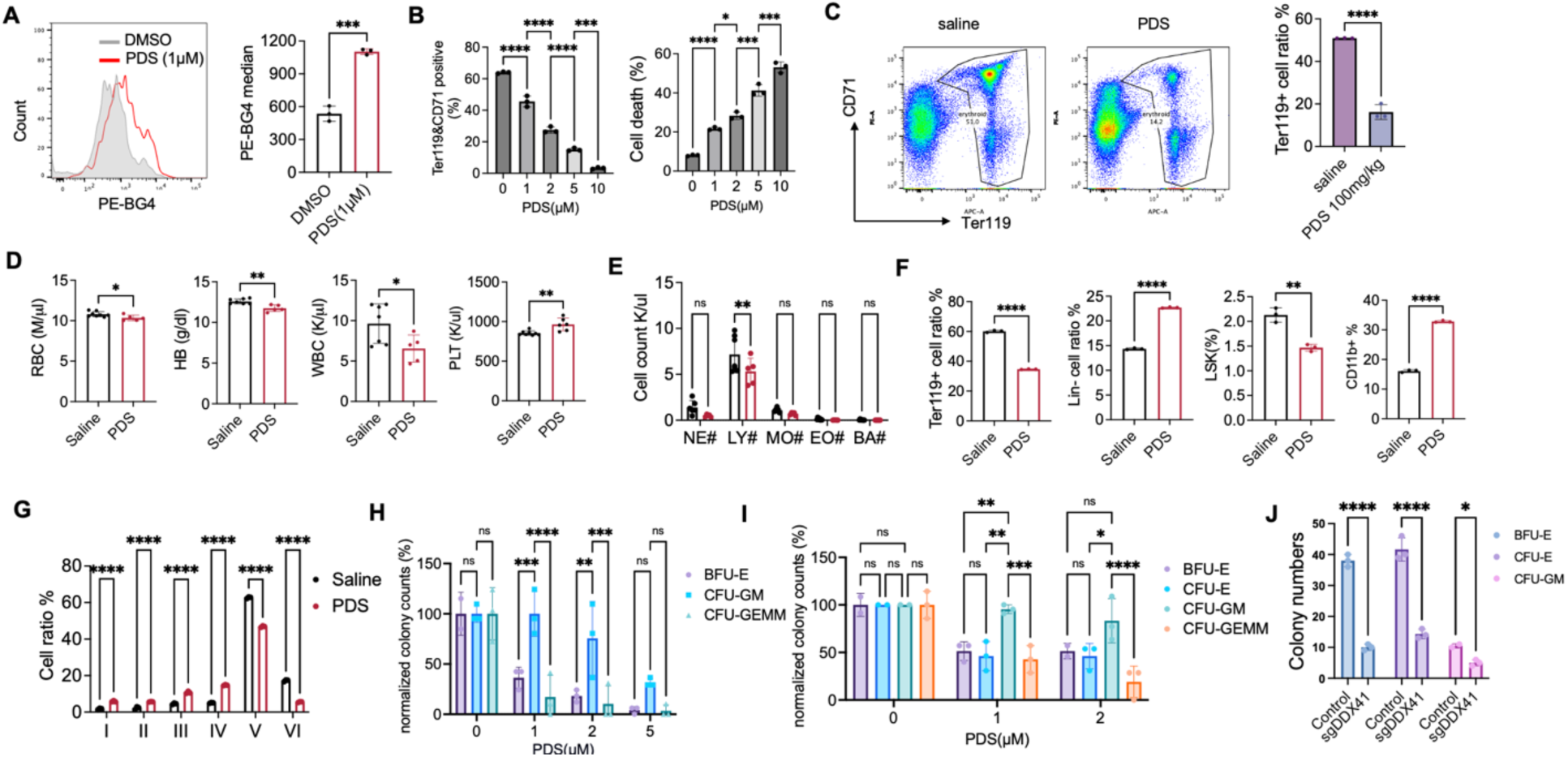
G4 accumulation is detrimental to erythropoiesis. (A) Bone marrow lineage-negative cells from 2-month-old wild-type mice were cultured in an Epo-containing medium for 24 hours, followed by 1 μM PDS treatment for 6 hours. The cells were then analyzed by flow cytometry using BG4 antibody. The quantification of flow cytometry results is on the right. (B) Bone marrow lineage-negative cells from 2-month-old wild-type mice were cultured in an Epo-containing medium with the indicated concentration of PDS. Cells were analyzed for differentiation and cell death at 24 hours by flow cytometry. The dead cells are defined as propidium iodide and annexin V double positive. (C) Two-month-old wild-type mice were treated with daily intraperitoneal injections of PDS at a dose of 100 mg/kg/day for three consecutive days. The mice were euthanized on day 3 for bone marrow examination of erythroid differentiation by flow cytometry. Quantification is on the right. (D) Two-month-old wild-type mice were chronically treated with PDS or saline. A complete blood count of the mice was performed two months following treatment. (E) Leukocyte count of D. (F) Quantification of flow cytometry of different lineages and HSPCs in mice from D. (G) Erythroid cell analysis with CD44 and forward scatter in mice from D. (H) Bone marrow lineage-negative cells from 2-month-old wild-type mice were cultured in MethoCult M3434 with the indicated concentration of PDS. Indicated colonies were quantified after 10 days. BFU-E: Burst-forming unit-erythrocyte; CFU-GM: Colony-forming unit-granulocyte-macrophage; CFU-GEMM: Colony-forming unit-granulocyte, erythrocyte, monocyte, and macrophage. The colony numbers are normalized to the untreated group. (I) Human CD34+ cells were cultured in MethoCult H4435 enriched medium with the indicated concentration of PDS. Colonies were quantified after 14 days. CFU-E: Colony-forming unit-erythrocyte. (J) The same colony assay as I except that human CD34+ cells were transduced with Cas9 and DDX41 sgRNA. Control: CRISPR V2 vector with scrambled sgRNA. All the error bars represent the SEM of the mean. The comparison between two groups was evaluated with 2 tailed t tests, and the comparison among multiple groups was evaluated with 1-way ANOVA tests. * p<0.05, **p<0.01, ***p<0.001, and ****p<0.0001. ns: not significant.

We performed murine in vitro colony-forming unit (CFU) assays under different concentrations of PDS. We found that erythroid-related colonies, such as BFU-E and CFU-GEMM, decreased more significantly than myeloid cell colonies (CFU-GM) in both colony number and size (**Figure 3H**). The same results were obtained when we performed CFU assays in human CD34+ cells (**Figure 3I**). We also knocked out DDX41 using CRISPR/Cas9 in CD34+ cells. We found significantly reduced BFU-E and CFU-E colonies (**Figure 3J**), indicating DDX41 could also be involved in stem cell commitment to the erythroid lineage. These results show that the erythroid lineage is more sensitive to G4 stresses, reinforcing our finding that DDX41 deficiency significantly impairs erythropoiesis.

### DDX41 is distributed at overlapping genomic loci as G4 and dissolves G4

The genome distribution of G4 has been studied in different cell types and various species^33–35^. The genome-wide G4 localization in hematopoietic cells and whether DDX41 co-localizes with G4 are unclear. To understand this, we performed CUT&RUN assays in purified maturing mouse bone marrow Ter119+ erythroblasts and bone marrow lineage-negative HSPCs. We found a significant colocalization of G4 and Ddx41 at the genome level both in HSPCs and erythroblasts. Their distributions are mainly in the intergenic and intron regions, consistent with a recent report^36^. Interestingly, G4 and Ddx41 genome occupation and colocalization are markedly increased in erythroblasts compared to HSPCs, consistent with the critical role of Ddx41 in erythropoiesis (**Figure 4A and 4B**). Motif analyses demonstrated that these G4 and Ddx41 enriched genome regions include binding loci of critical transcription factors for erythropoiesis, such as Tal1 and Gata1 (**Supplemental Figure 6A**). We further confirmed these genomic studies using a confocal immunofluorescence assay of G4 and Ddx41 in cultured mouse erythroblasts, which indeed showed their co-localization (**Figure 4C**). Computational predictions indicate that ribosomal DNAs (rDNAs) are highly enriched for putative quadruplex formation due to their repetitive DNA sequences^33,37^. Consistent with these reports, we found enrichment of G4 and Ddx41 on rDNAs in both HSPCs and erythroid cells (**Supplemental Figure 6B and 6C**).

**Figure 4.**
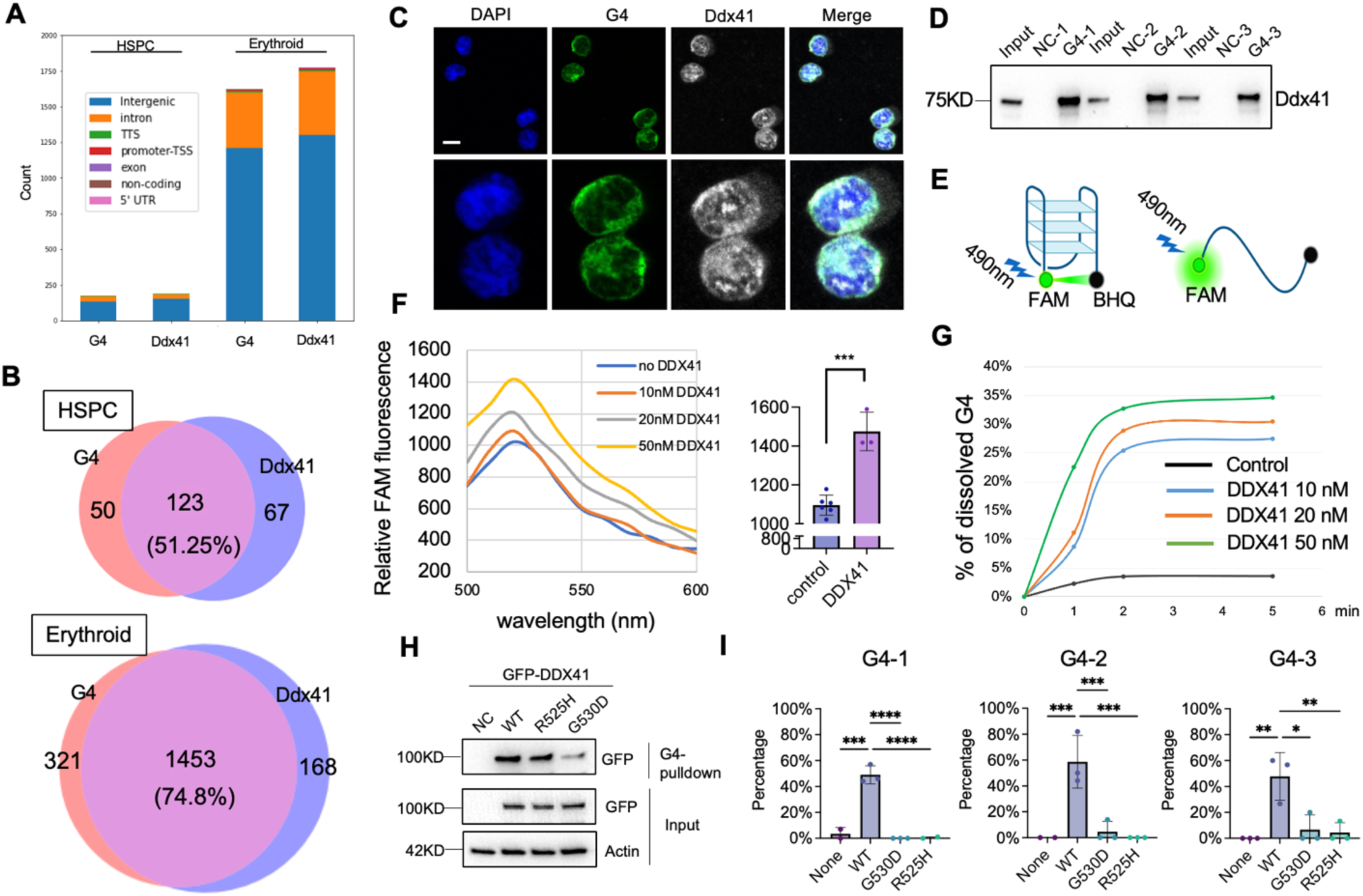
DDX41 binds to and dissolves G-quadruplexes. (A) Genome-wide distributions of G4 and Ddx41 binding sites in mouse lineage negative HSPCs and Ter119 positive erythroid cells through CUT&RUN assays. (B) Quantification of the co-localized genomic loci from A. (C) Immunofluorescence analyses of G4 and Ddx41 in mouse lineage-negative cells cultured in Epo medium for 1 day. High-magnification images of the bottom two cells are shown in the lower panels. Scale bar: 5 μm. The pictures are representative of >10 different fields. (D) Mouse bone marrow lineage-negative cells were cultured in an Epo-containing medium for 1 day. The lysates were incubated with the indicated G4 or non-G4 control (NC) oligonucleotides in the presence of KCl for 1 hour. Western blotting of Ddx41 was then performed. (E) Schematic illustration of the FRET assay. (F) Dose-dependent increase of FAM signal with increased concentration of recombinant human DDX41 protein. Statistical analysis of FAM signals with 50 nM DDX41 compared to the control group is on the right. (G) Time course of DDX41-mediated G4 dissolving activity. (H) Western blotting analyses of indicated proteins in the G4 pull-down assay using biotin-conjugated G4 or non-G4 sequences and lysates from HEK293T cells that were transfected with GFP-tagged wild-type or mutant DDX41. (I) FRET assays using G4s as in F incubated with wild-type or mutant recombinant DDX41. The Y-axis represents the percentage of unfolded G4 structures as in G. All the error bars represent the SEM of the mean. The comparison between two groups was evaluated with 2 tailed t tests, and the comparison among multiple groups was evaluated with 1-way ANOVA tests. * p<0.05, **p<0.01, ***p<0.001, and ****p<0.0001.

The co-occupation of DDX41 and G4 in the erythroid genome and the upregulation of G4 after DDX41 depletion indicate that DDX41 functionally maintains the level of G4. DEAD box family proteins, such as DDX5, were known to resolve DNA G4^9,17^. To determine whether DDX41 dissolves G4, we first performed a pull-down assay using biotin-conjugated G4 oligos. The canonical G4 motif is Gm-Xn-Gm-Xo-Gm-Xp-Gm where each G-tract (Gm, m=2-4) is separated by loops (Xn, Xo, and Xp), and n, o, and p are the combination of nucleotides of various lengths (up to 7) ^28^. We designed three different G4 oligonucleotides. Three non-G4 oligonucleotides were used as controls. These oligonucleotides were annealed with biotinylated counterparts and then captured on streptavidin magnetic beads. They were subsequently incubated with lysates from cultured mouse erythroblasts in the presence of K^+^ cations. DDX41 exhibited specific binding to all three G4 structures of distinct topologies (**Figure 4D**), indicating its ability to recognize a wide range of G4 structures.

To determine whether DDX41 directly dissolves G4, we performed an in vitro fluorescence resonance energy transfer (FRET) assay in which a fragment of G4 DNA is flanked by 6-fluorescein (6-FAM) on the 3’-end and black hole-1 quencher on the 5’-end^17^ (**Figure 4E**). Recombinant human DDX41 was added in vitro, together with G4-FRET and K^+^ cations. Indeed, we found a dose-dependent increase of the 6-FAM fluorescence with the increasing amount of DDX41, demonstrating that G4 was dissolved by DDX41 (**Figure 4F, Supplemental Figure 7A**). The kinetics of DDX41 in dissolving G4 was also rapid and dose-dependent (**Figure 4G, Supplemental Figure 7B**). We next tested two of the most common somatic mutations of *DDX41*, R525H, and G530D on their influences on G4 binding capacities. We found that R525H mildly compromised the binding, whereas G530D markedly reduced it (**Figure 4H**). We further discovered that both mutants lost G4 dissolving activity in all three G4s we tested (**Figure 4I**). These data establish DDX41 as a G4 dissolver, which is compromised by its loss of function mutations.

### DDX41 deficiency-mediated G4 accumulation leads to genome instability, ribosomal biogenesis defects, and upregulation of p53

Accumulation of G4 is associated with genome instability and defects in ribosome biogenesis due to G4 enrichment on rDNAs^33,37–41^. Compromised ribosome biogenesis during erythropoiesis is well known to trigger p53-mediated cell death^42^, which is believed to be the pathogenesis of Diamond-Blackfan anemia (DBA) and contribute to the development of del(5q) MDS^43,44^. We found increased γ-H2AX (a marker for genome instability) when the cultured mouse bone marrow HSPCs were treated with PDS (**Figure 5A-5C**). As expected, the transcription of ribosomal RNAs was significantly reduced (**Figure 5D**). Defects in ribosomal RNA biogenesis are known to negatively influence the expression of ribosomal proteins^43,45^. Consistently, we found PDS treatment significantly reduced many ribosomal proteins, including Rps19 (mutated in 25% of DBA patients), Rps14 (deleted in del(5q) MDS patients), and Rpl26 (mutated in certain DBA patients)^45^. The level of p53 was also increased (**Figure 5E**). To directly investigate how DDX41 deficiency affects ribosomal biogenesis in vivo, we used *HBBCre:Ddx41^fl/fl^* mice since these mice survive and Ddx41 is depleted in the late-stage erythroblasts when the cells are most abundant. We cultured the bone marrow lineage negative HSPCs from these mice in Epo-medium and found significantly decreased transcription of rRNAs on day 1 when Ddx41 starts to reduce, and G4 is significantly increased (**Figure 5F and 2I**). The ribosomal protein levels remain steady on day 1, possibly due to the stability of proteins compared to RNAs, but significantly reduced on day 2 (**Figure 5G**). Interestingly, we found no increase in γ-H2AX when Ter119-negative erythroblasts from *HBBCre:Ddx41^fl/fl^*or *EpoRCre:Ddx41^fl/+^* mice were cultured in vitro (**Supplemental Figure 7C**). These results indicate that these lineage-negative cells may be adapted to the Ddx41 deficiency during development.

**Figure 5.**
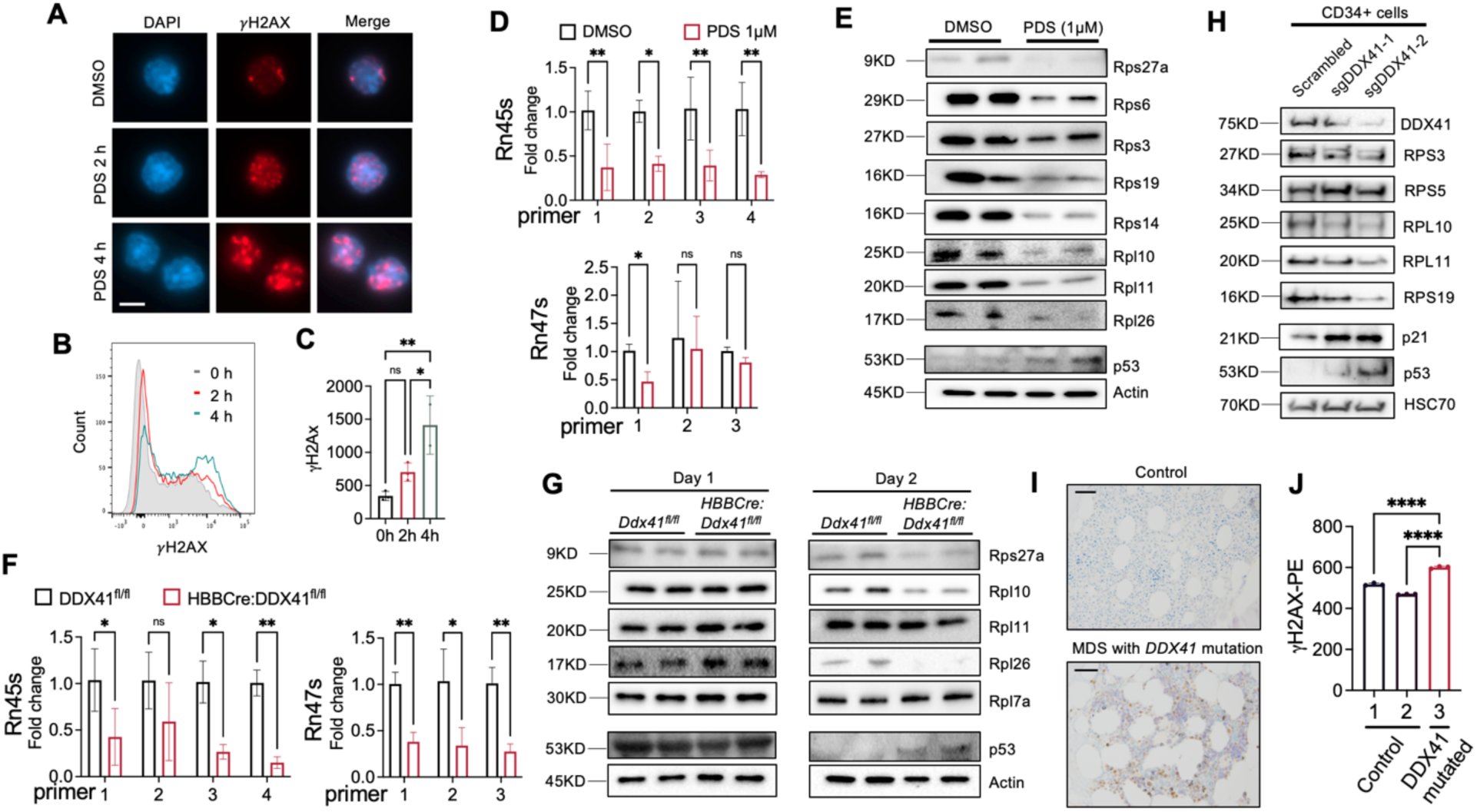
DDX41 deficiency-mediated G4 accumulation leads to genome instability, ribosomal biogenesis defects, and upregulation of p53. (A) Epo medium-cultured mouse bone marrow lineage negative HSPCs were treated with 1 μM PDS for the indicated time. Immunofluorescence assays of γ-H2AX were performed, and representative images of the erythroid cells were presented. Scale bar: 5 μm. (B) Flow cytometry assay of the cells in A. (C) Statistical quantification of γH2AX signals in B. (D) Epo medium-cultured mouse bone marrow lineage negative HSPCs were cultured for 1 day, followed by the treatment of 1 μM PDS for 6 hours. Quantitative RT-PCR analyses of indicated ribosome RNAs were performed using different primer sets. (E) Western blotting assays of indicated in cells from D. Actin was used as a loading control. (F) Same as D except that bone marrow lineage negative HSPCs from *HBBCre:Ddx41^fl/fl^*mouse were cultured for 1 day before the quantitative RT-PCR assays. (G) Western blotting assays of the indicated proteins in F. Cells from both day 1 and day 2 cultured cells were analyzed. (H) CD34+ cells were transduced with lentiviral vectors expressing indicated sgRNAs and Cas9. Cells were then harvested for Western blotting of the indicated proteins at day 9 in culture. (I) Immunohistochemical stains of p53 in bone marrow core biopsies from the patient in Figure 1J and a normal individual. Scale bar: 100 μm. (J) Quantification of γ-H2AX in bone marrow mononuclear cells from the patient in I and 2 control individuals. All the error bars represent the SEM of the mean. The comparison between two groups was evaluated with 2 tailed t tests, and the comparison among multiple groups was evaluated with 1-way ANOVA tests. * p<0.05, **p<0.01, ns: not significant.

Consistent with these findings in mouse erythroid cells, DDX41 knockout in CD34+ HSPCs led to a similar reduction of various ribosomal proteins and upregulation of p53. The p53 downstream target p21 was also increased (**Figure 5H**). We next tested whether the p53 level increased in the patient with *DDX41* mutation. We used bone marrow biopsies from the same patient with the G530D mutation and performed an immunohistochemical stain for p53. We found that the p53 level (**Figure 5I**) and γ-H2AX (**Figure 5J**) significantly increased in the patient with *DDX41* mutation compared to the normal control, consistent with the findings in the mouse models.

### Activation of the cGAS-STING pathway, but not p53 upregulation, is essential for DDX41 deficiency-mediated ineffective erythropoiesis

While p53 mediates many pathologies of red cell-related diseases, it has been documented that overexpression of p53 does not induce overt abnormalities in a transgenic model ^46^. To determine whether p53 mediates the major phenotypes of Ddx41 hematopoietic specific deficiency mice as it does in DBA and del(5q) MDS, we took a genetic approach and crossed p53 knockout mice with *VavCre:Ddx41^fl/+^* mice. If p53 is critical, p53 deficiency would rescue the lethality of *VavCre:Ddx41^fl/fl^* mice. However, no surviving *VavCre:Ddx41^fl/fl^ Trp53*^-/-^ mice were born (**Figure 6A and 6B**). Dissection of the pregnant mice revealed that *VavCre:Ddx41^fl/fl^ Trp53*^-/-^ embryos remained pale with an underdeveloped fetal liver (**Figure 6A**). These results demonstrate that p53 is not essential to mediate ineffective erythropoiesis and cell death in the Ddx41 hematopoietic-specific knockout mouse model.

**Figure 6.**
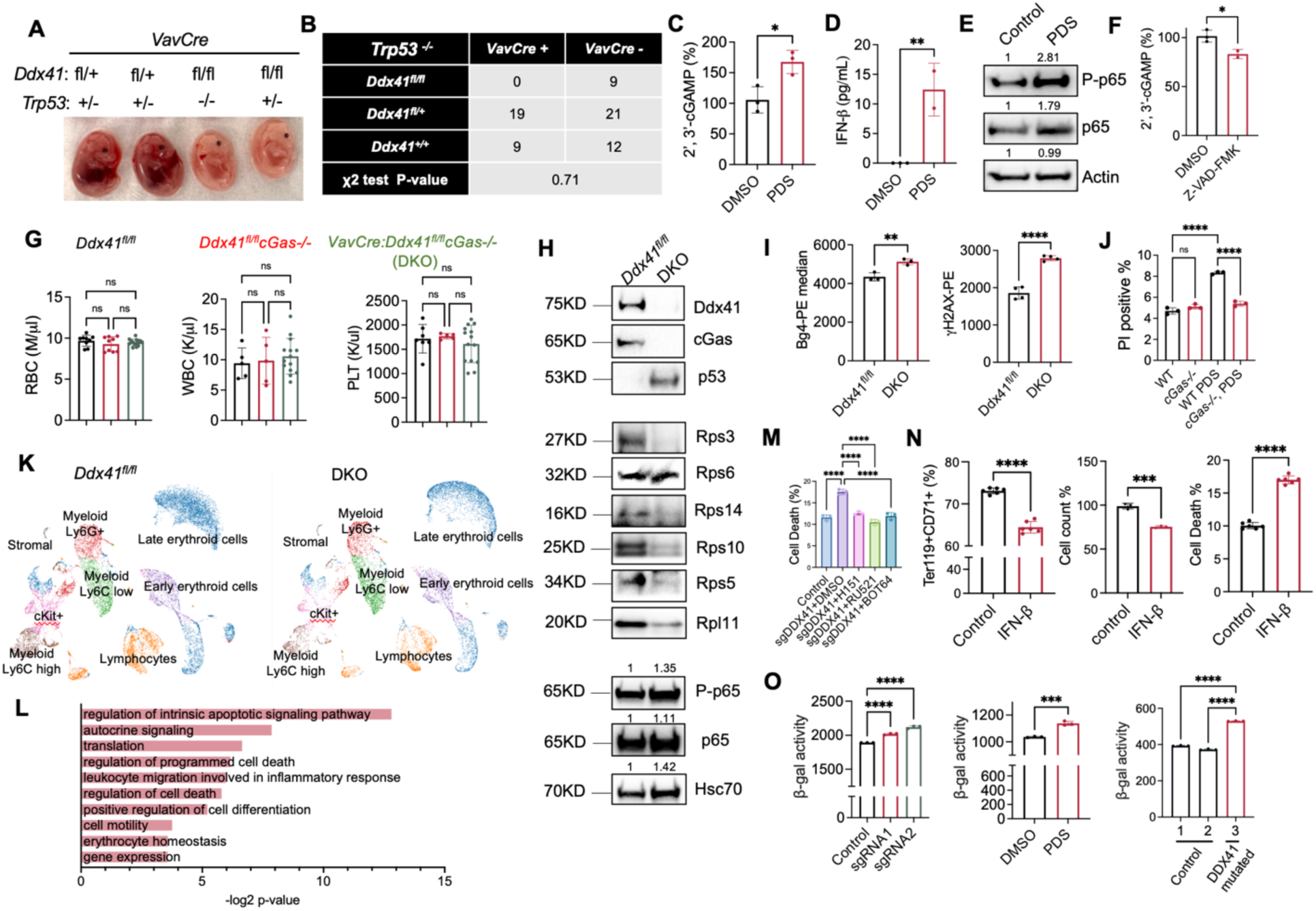
Activation of the cGAS-STING pathway, but not p53 upregulation, is essential for DDX41 deficiency-mediated ineffective erythropoiesis. (A) Representative pictures of E14.5 embryos with the indicated genotypes. (B) Mendelian segregation of indicated mice. A χ^2^ test yields a p-value of 0.71, indicating no significant deviation from the expected Mendelian segregation. (C-D) Mouse lineage-negative HSPCs were cultured in Epo medium for 1 day, followed by treatment with 1 μM PDS for 6 hours. Cells were collected for 2’,3’-cGAMP (C) and IFN-β (D) ELISA assays. (E) Western blotting of indicated proteins in cells from C. The bands were also quantified for phospho-p65, total p65, and actin. (F) Mouse lineage-negative HSPCs were cultured in Epo medium for 1 day, followed by treatment with 40 μM Z-VAD-FMK for 6 hours. Cells were then collected for ELISA analysis of 2’,3’-cGAMP. (G) Complete blood count of indicated mice at 2 months old. (H) Western blotting assays of indicated proteins from the bone marrow Ter119+ erythroid cells of mice from G. The bands were also quantified for phospho-p65, total p65, and Hsc70. (I) G4 and γH2AX levels were analyzed by flow cytometry and quantified from the bone marrow Ter119+ erythroid cells in mice from G. (J) Lineage-negative cells from the indicated mice were cultured in Epo medium for 1 day followed by treatment with 1 μM PDS for 6 hours. Cell death was analyzed by flow cytometry. (K) Uniform Manifold Approximation and Project (UMAP) plots showing the distribution of annotated populations in the bone marrow of indicated mice at 2 months old. (L) Pathway enrichment analyses of down-regulated genes in the late erythroid cell population in DKO mice in K. (M) CD34+ cells were cultured in Epo medium for 4 days, followed by DDX41 knockout using the CRISPR/Cas9 and treatment of the indicated inhibitors simultaneously for 2 days. Flow cytometry was utilized to analyze cell death post-treatment. The control group is cells treated with a CRISPR vector containing scrambled sgRNA. The dead cells are defined as propidium iodide and annexin V double positive. (N) Mouse lineage-negative cells were cultured in Epo medium for 1 day with or without 100 ng/ml IFN-β, followed by flow cytometry analyses of cell differentiation, proliferation, and cell death. The dead cells are defined as propidium iodide and annexin V double positive. (O) Quantitative analyses of β-galactosidase activity in cells from CD34+ cells with DDX41 knockout, mouse lineage negative cells treated with 1 µM PDS for 24 hours in culture, and the bone marrow mononuclear cells from the MDS patient in Figure 5J. CD34+ cells were transduced with DDX41 sgRNA on day 6 of culture, and β-galactosidase activity was determined 24 hours after transduction. All the error bars represent the SEM of the mean. The comparison between two groups was evaluated with 2 tailed t tests, and the comparison among multiple groups was evaluated with 1-way ANOVA tests. ns: not significant. * p<0.05, **p<0.01, ***p<0.001, and ****p<0.0001.

DDX41 was reported to sense intracellular dsDNA and activate STING to mediate type I interferon response in dendritic cells ^11,47^. Indeed, we found the level of cGAMP, intracellular second messenger in response to cGAS activation, was significantly increased in PDS-treated erythroid cells (**Figure 6C**). The downstream targets of the cGAS-STING pathway, including interferon beta (IFN-β) and the NF-κB signaling, were also activated (**Figure 6D and 6E**). We previously revealed that erythroid cells generate transient nuclear openings in the early stages of terminal erythropoiesis^48,49^, which could further activate cGAS. To test this, we treated the cultured mouse bone marrow erythroid cells with a caspase inhibitor, which blocks nuclear opening. This led to a significant reduction of cGAMP (**Figure 6F**), suggesting that nuclear openings contribute to the vulnerability of the erythroid cells to the genome instability induced by G4 upregulation.

We then took a similar genetic approach and crossed cGas knockout mice with *VavCre:Ddx41^fl/+^* mice. Intriguingly, the cGas deficiency completely rescued the embryonic lethality of the *VavCre:Ddx41^fl/fl^* mice. *VavCre:Ddx41^fl/fl^cGas-/-* (DKO) mice showed no evidence of anemia or other cytopenias (**Figure 6G, Supplemental Figure 8A, and 8B**). The bone marrow hematopoiesis was also intact (**Supplemental Figure 8C**). Western blotting of the bone marrow Ter119+ erythroid cells confirmed cGas and Ddx41 deletions but also showed loss of ribosomal proteins and upregulation of p53 in the DKO mice (**Figure 6H**), which indicates that G4 upregulation-mediated genome instability leads to parallel activation of the cGas and p53 pathways. Consistent with this indication, the level of phospho-p65 was unchanged (**Figure 6H**), whereas G4 and γ-H2AX remained upregulated in the bone marrow Ter119+ erythroid cells of the DKO mice compared to the cells in the control mice (**Figure 6I**). The critical role of cGas in mediating Ddx41 deficiency-induced pathogenesis was further evidenced by the resistance of primary erythroid cells from cGas knockout mice to PDS-mediated cell death (**Figure 6J**). We next performed a single-cell RNA sequencing assay of the total bone marrow cells from DKO mice and their littermate control (**Supplemental Figure 8D**). While non-erythroid hematopoietic populations showed no apparent distinctions, late erythroid cells in DKO mice exhibited altered gene expression, including down-regulation of genes involved in cell death regulation and erythroid hemostasis (**Figure 6K, 6L, and Supplemental Figure 8E, Supplemental Table 2**).

cGAS was shown to translocate to the nucleus under DNA double-strand break to suppress DNA repair independent of STING^50^. We found that cGas, as well as Sting, was predominantly located in the cytoplasm in the erythroid cells upon PDS treatment, demonstrating a cGas-Sting-dependent pathway and consistent with the upregulation of IFN-β and the activation of NF-κB signaling (**Supplemental Figure 8F**). In line with the role of the cGAS-STING pathway, treatment of DDX41 deficient CD34+ cells with a cGAS inhibitor (RU521), a STING inhibitor (H151), or an NF-κB inhibitor (BOT64) significantly rescued cell death (**Figure 6M**). Similar to PDS, treatment of the erythroid cells with IFN-β significantly compromised cell differentiation, proliferation, and induced cell death (**Figure 6N**).

These data point towards cGAS-STING activation-induced senescence, instead of p53-induced cell death, in mediating ineffective erythropoiesis in DDX41 deficiency. Consistently, we observed increased senescence-associated β-galactosidase activities in DDX41 CRISPR knockout or PDS-treated CD34+ cells in Epo-containing medium, as well as bone marrow mononuclear cells from the *DDX41* mutated patient (**Figure 6O**).

### Human erythroid cells are sensitive to DDX41 deficiency in an iPSC-derived bone marrow organoid environment

To extend these findings to the human bone marrow in vivo setting, we established an induced pluripotent stem cell (iPSC)-derived human bone marrow organoid system resembling primary human bone marrow biopsy samples (**Figure 7A**). Whole-mount 3D imaging revealed an endothelial network accompanied by stromal cells along the capillary wall and hematopoietic cells arranged in clusters within and outside the vessels (**Figure 7B**). Erythropoiesis in the organoid mirrors primary human bone marrow samples, forming tight erythroid islands (**Figure 7C**). A flow cytometry assay further confirmed multilineage hematopoiesis in the organoids (**Figure 7D**). To determine whether these organoids can be applied to study human ex vivo bone marrow engraftment, we incubated the organoids with CellVue-labeled donor bone marrow CD34+ cells for three days. The donor cells could be readily detected and surrounded by the recipient hematopoietic cells in the organoids (**Figure 7E**). These donor cells could also differentiate into mature hematopoietic cells detected by immunofluorescence and flow cytometry assays (**Figure 7E and 7F**). With this system, we depleted DDX41 through CRISPR in CD34+ cells and incubated these cells with the bone marrow organoids. As expected, the CD71+ erythroid cells, but not CD11b+ myeloid cells, derived from the donor CD34+ cells transduced with DDX41 sgRNA were significantly reduced compared to the control donor cells. Interestingly, the erythroid cells derived from the recipient bone marrow organoids were also significantly reduced, indicating a potential indirect impact from the inflammatory environment due to the cGAS-STING activation (**Figure 7G**).

**Figure 7.**
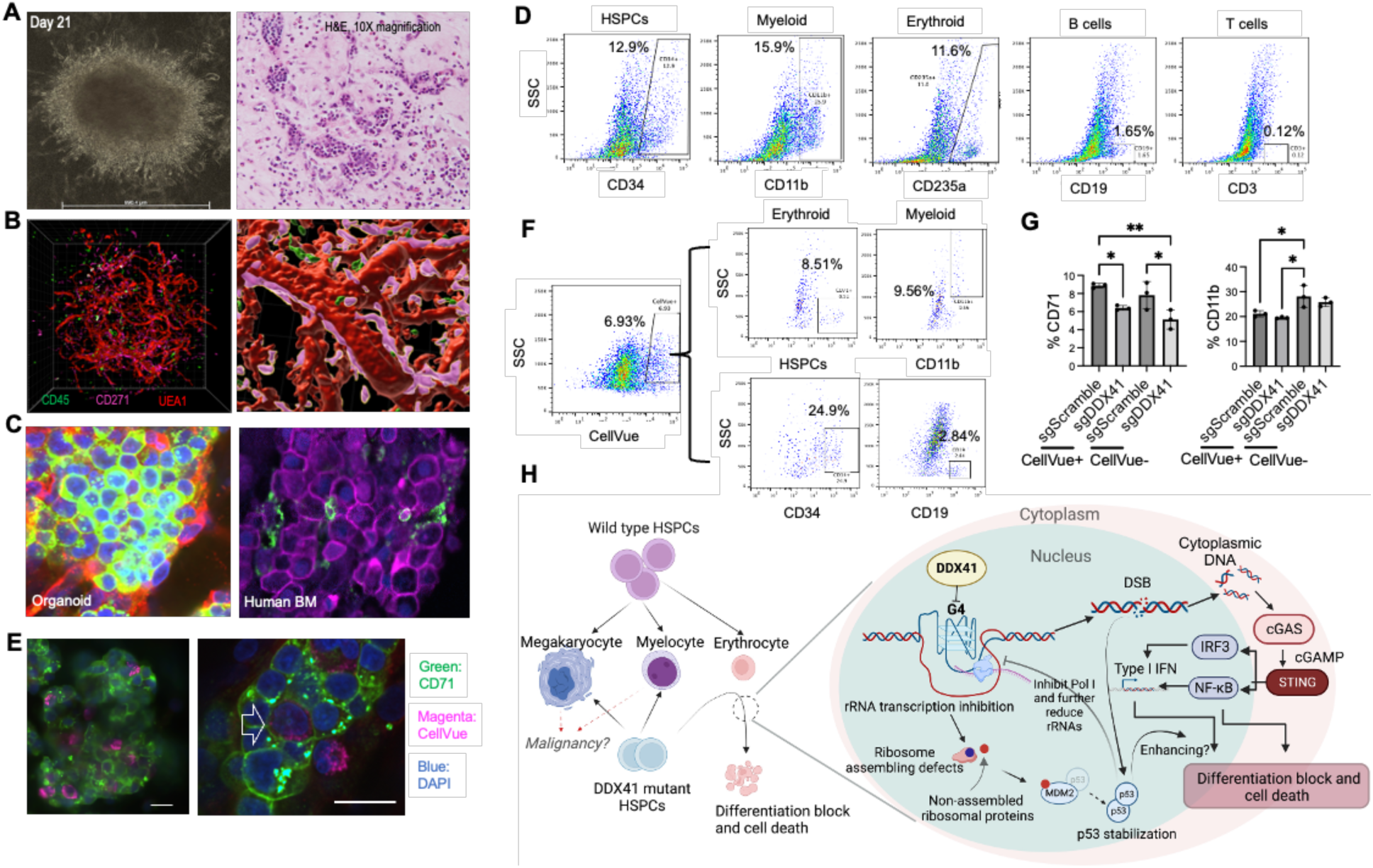
Human erythroid cells are sensitive to DDX41 deficiency in an iPSC-derived bone marrow organoid environment. (A) Representative wide-field picture and H&E stains of bone marrow organoid in culture. (B) Whole-mount 3D imaging of the organoids. Imaris was used for cell surface rendering. Organoids were stained with indicated antibodies and subsequently imaged using a laser scanning confocal platform. (C) Confocal immunofluorescence assays of erythroid islands in the iPSC-derived bone marrow organoids (left) and a primary human bone marrow biopsy (right). CD71 was labeled with green for organoids and magenta for primary bone marrow. DAPI: blue. (D) Flow cytometry assays of the organoids using indicated antibodies for various lineages. (E) 10,000 CellVue-labeled donor CD34+ HSPCs were co-incubated with iPSC-derived bone marrow organoids for 3 days in each well of a 96-well plate, followed by an immunofluorescence assay. Representative pictures show the engraftment of donor hematopoietic cells into the organoid. Green, red, and blue represent CD71, CellVue, and DAPI-positive nuclei, respectively. The arrow points to an engrafted CellVue positive cell expressing CD71. (F) Flow cytometry of the organoids using indicated antibodies for various lineages of the engrafted cells in organoids from E. (G) Same as E, except the donor CD34+ cells were transduced with lentiviral vectors expressing Cas9 and indicated sgRNAs before co-incubation. After 3 days, the cells were collected for flow cytometric assays of erythroid and myeloid differentiation of CellVue-positive donor hematopoietic cells and negative iPSC-derived hematopoietic cells. Each data point represents cells combined from 10 organoids. The comparison was evaluated with 1-way ANOVA tests. * p<0.05, **p<0.01. (H) Schematic model of the function of DDX41 during erythropoiesis. The diagram is generated through BioRender.

We also treated the organoid with PDS for 24 hours and analyzed erythropoiesis using flow cytometry. Since the organoid already contains erythroid cells at various stages of development, this approach allows us to study the responses of cells at different stages of erythropoiesis to G4 accumulation. Consistent with the results from other experiments, we found a significant decrease in the population of CD34+CD71+ cells, which include mainly early-stage erythroid cells. However, no significant differences were found in the population of CD34-, CD71+, and CD235a+ cells, which represent late-stage erythroid cells (**Supplemental Figure 8G**).

## Discussion

Our study shows that DDX41 directly interacts with and resolves G-quadruplexes. DDX41 deficiency increases G4 levels, causing genome instability and impairing ribosomal biogenesis. This elevation in G4s activates p53 and the cGAS-STING pathway. Genetic studies in mice confirm that knocking out cGas, but not p53, rescues the lethality in hematopoietic-specific Ddx41 knockout mice (**Figure 7H**).

In *VavCre:Ddx41^fl/fl^ cGas-/-* (DKO) mice, the sustained reduction in ribosomal proteins and increased p53 levels mirror mechanisms seen in ribosomopathies like Diamond-Blackfan Anemia (DBA). In DBA, impaired ribosome assembly results in excess free ribosomal proteins, which bind to MDM2, reducing its ability to degrade p53^45^. Despite ribosomopathies, the complete rescue of survival in the DKO mice is consistent with reports that an increase in p53 activity or haploinsufficiency of *MDM2* does not necessarily result in severe defects in mice ^46,51–54^. Nevertheless, ribosomopathies and the continued presence of genome instability due to G4 upregulation increase the possibility of the risk of the development of hematologic malignancies in these mice, especially with aging. It is also possible that the phenotypically normal myeloid, monocytic, and dendritic-specific Ddx41 knockout mice could develop hematologic diseases with aging due to the accumulation of DNA double-strand break with the upregulation of G4. The increased G4, γ-H2AX, and p53 levels in the MN patient with somatic DDX41 mutation are consistent with this possibility.

Our study reveals erythroid cells’ hypersensitivity to DDX41 deficiency, likely due to widespread G4 structure enrichment, especially on ribosomal DNAs. Erythroid cells are vulnerable to replication stress from DNA secondary structures like G4, explaining the detrimental effects of DDX41 deficiency or PDS treatment on their differentiation and survival. Similar sensitivity of erythroid precursors to Ddx41 deficiency was also reported in zebrafish^26^, highlighting the essential roles of DDX41 in governing erythroid genome stability across species. Under physiologic conditions, ribosome biogenesis is critical in the early stages of erythropoiesis but decreases in the late stages of terminal erythropoiesis by the decline in rDNA transcription^55^. This is in line with our findings that there are no significant phenotypes in *HBBCre:Ddx41^fl/fl^* mice despite an increase in G4 since G4 upregulation-mediated defects occur in the late stages of terminal erythropoiesis when the ribosomal biogenesis is naturally reduced, and chromatin is markedly condensed^31,48,56^. It is also possible that the activation of the cGAS-STING pathway due to DDX41 deficiency is also not essential at the late stages of terminal erythropoiesis.

These findings in murine models were confirmed in human settings using primary patient samples and iPSC-derived bone marrow organoids. The somatic G530D mutation identified in our patient is typically absent in the erythroid population, aligning with DDX41 loss-of-function effects during erythropoiesis, particularly with aging. Similar impairments were observed when DDX41 knockout HSPCs were engrafted into iPSC-derived organoids, leading to compromised erythroid maturation and senescence. Notably, these effects were also evident in erythroid cells from the organoids, consistent with the activation of the cGAS-STING pathway in the inflammatory bone marrow environment, which renders erythroid cells particularly sensitive^57^.

## Materials and methods

### Mice

Wild-type C57BL/6J mice (strain #000664), cGas knockout mice (strain #026554), Cre-recombinase expression mice Vav-Cre (strain #008610), EpoR-Cre (strain #035702), MRP8-Cre (strain #021614), CD11c-Cre (strain #008068), and LysM-Cre (strain #004781) were purchase from the Jackson Laboratory. HBB-Cre was provided by Nicolae Valentin David. DDX41 floxed (Ddx41^fl/fl^) mice were provided by Susan Ross.

Tissue-specific Ddx41 knockout mice were generated by breeding tissue-specific Cre recombinase expression mice with Ddx41^fl/fl^ mice. The F1 generation, which carried Cre recombinase and was Ddx41^fl/+^, was subsequently crossed with Ddx41^fl/fl^ mice. For Ddx41^fl/fl^ breeding with Vav-Cre, female Vav-Cre-carrying mice were bred with male Ddx41^fl/fl^ mice to avoid Ddx41 knockout during spermatogenesis.

All animal studies followed the Guidelines for the Care and Use of Laboratory Animals and were approved by the Institutional Animal Care and Use Committees at Northwestern University.

### Mouse bone marrow and spleen flow cytometry assay

Mice were euthanized following ethical guidelines approved by the IACUC at Northwestern University. Femur bones were dissected to extract bone marrow cells via PBS flushing. Spleens were weighed and subsequently homogenized. These isolated single cells were labeled with antibodies for subsequent flow cytometric analysis. For erythroid cell characterization, APC-Ter119, PE-CD71, and FITC-CD44 antibodies were employed. Myeloid cell analysis utilized APC-CD11b, BV421-ly6C, and PE-ly6G antibodies, while lymphoid cell populations were assessed using PE-B220 and APC-CD3 antibodies. For HSPC analysis, the cells were stained with BD Pharmingen™ PerCP-Cy™5.5 Mouse Lineage Antibody Cocktail along with FITC-Ly5A/E and APC-CD117. Subsequently, cells that exhibited a negative signal in the PerCP-Cy5.5 channel were selected, and further analysis was conducted on these cells using the FITC and APC channels to identify and characterize LSK (Lin-Sca1+Kit+) cells. For cell death analysis, cells were stained using the BD Pharmingen™ FITC Annexin V Apoptosis Detection Kit I and subsequently analyzed via flow cytometry.

### Quantification of G-quadruplexes using flow cytometry

The quantification of G-quadruplexes was performed using the BG4 antibody with mouse IgG1 Isotype. Cells were fixed in 0.25% Glutaraldehyde/PBS for 1 hour, followed by permeabilization with 0.1% Triton-X/PBS. To remove the RNAs, cells were incubated in 0.05% Triton-X/PBS with 3% bovine serum albumin (BSA) and 10 µg/ml RNaseA for 30 minutes. Subsequently, they were incubated in PBS containing 0.05% Triton-X, 3% BSA, and 10 µg/ml BG4 antibody for at least 30 minutes. After washing, cells were resuspended in PBS with PE-conjugated anti-mouse IgG1 antibody for 10 minutes. Finally, the cells were washed and resuspended in PBS for flow cytometry analyses. Median fluorescence intensity was used for statistical analyses to mitigate the influence of outliers and skewed distributions.

### Antibodies

The following antibodies (vendor, catalog number) were used for Western blotting assays: DDX41 (CST, #15076), BG4 (Absolute Antibody, Ab00174), p53 (CST, #9282), cGAS (E5V3W) (CST, #79978), Phospho-Histone H2A.X (Ser139) (D7T2V) (CST, #80312), NF-κB Pathway Antibody Sampler Kit (CST #9936), RPL26 (Proteintech, #17619-1-AP), RPL7A (Proteintech, 15340-1-AP), RPS27A (Proteintech, #14946-1-AP), RPS3 (Proteintech, #11990-1-AP), RPS6 (Proteintech, #14823-1-AP), RPS19 (Proteintech, #15085-1-AP), RPS14 (Proteintech, 16683-1-AP), Ribosomal Protein S3 (D50G7) (CST, #9538S), S6 Ribosomal Protein (5G10) (CST, #2217), RPL10 (Proteintech, #72912), RPL11 (D1P5N) (Proteintech, #18163), RPL5 (Proteintech, #14568), HRP-conjugated beta actin monoclonal antibody (Proteintech, #HRP-66009), anti-rabbit IgG, HRP-linked antibody (CST, #7074), and anti-mouse IgG, HRP-linked antibody (CST, #7076).

For flow cytometry assays, the following antibodies were used: APC rat anti-mouse TER119 (BD, #561033), FITC anti-mouse CD71 (Biolegend #113805), PE rat anti-mouse CD71 (BD, #567206), PE anti-mouse IgG1 (Biolegend, #406607), PE rat anti-mouse CD45R/B220 (BD, #553089), APC rat anti-mouse CD3 (BD, #565643), Phospho-histone H2A.X (Ser139) (CR55T33), PE F(ab’)2 fragment (Alexa Fluor® 488 conjugate) (CST, #4412), anti-mouse IgG (H+L), F(ab’)2 fragment (Alexa Fluor® 647 conjugate) (CST, #4410), Phospho-Histone H2A.X (Ser139) monoclonal antibody (Invitrogen, 12-9865-43), anti-rabbit IgG F(ab’)2 fragment (Alexa Fluor® 488 Conjugate) (CST, #4412), anti-mouse IgG (H+L) F(ab’)2 fragment (Alexa Fluor® 647 Conjugate) (CST, #4410), FITC rat anti-mouse CD44 (BD, #561859), APC rat anti-CD11b (BD, #553312), BV421 rat anti-mouse Ly-6C (BD, #562727), PE rat anti-mouse Ly-6G (BD, #561104), FITC rat anti-mouse Ly-6A/E (BD, #557405), and APC rat anti-mouse CD117 (BD, #553356).

### Isolation of bone marrow and fetal liver hematopoietic stem and progenitor cells (HSPCs)

Mouse femurs were dissected with surrounding tissues removed. The bone marrow was collected by flushing the bone cavities using 10 ml of phosphate buffer saline (PBS) supplemented with 2% fetal bovine serum (FBS). The harvested cells were then incubated for 10 minutes with a cocktail of biotin-conjugated lineage antibodies against CD3e, CD11b, CD45R, Ly-6G/C, and Ter119 (BD, #559971), followed by washing with PBS containing 2% FBS. Subsequently, the cells were incubated with streptavidin particles plus beads (BD, #557812) for 10 minutes and placed on a magnetic rack for 15 minutes for magnetic separation. The supernatant, enriched with lineage negative HSPCs, was collected for subsequent cell counting and analyses.

For fetal liver HSPC purification, E13.5 fetal liver was homogenized by repeated pipetting and filtering through a 40 μm cell strainer to obtain single cells. The filtered cells were incubated with biotin anti-mouse Ter119 antibody for 15 minutes and washed with PBS. After washing, the cells were resuspended, and Ter119-positive cells were pulled down with streptavidin magnetic beads. The remaining Ter119 negative HSPCs were then used for the follow-up experiments.

### In vitro erythroid differentiation of mouse lineage negative cells

The isolated lineage-negative cells were cultured in erythropoietin (Epo)-containing medium (IMDM supplemented with 15% FBS, 10% BSA, 10 μg/ml insulin, 200 μg/ml holo-transferrin, 0.1 mM β-mercaptoethanol, 1% penicillin-streptomycin, 2 mM glutamine, and 2 unit/ml Epo) at a density of 5 × 10^5^ cells/ml. The cells were then incubated for up to 48 hours at 37°C with 5% CO_2_. Cells were then harvested for analysis at different time points based on the experiments.

### Human bone marrow cells and Sanger sequencing

Total bone marrow cells from MDS patients were obtained following informed consent under institutional review board-approved protocols at Northwestern University. Isolation of specific cell populations was achieved using a bead-based method through biotin-conjugated CD34+ and CD71+ antibodies. Following isolation, the cells underwent a washing process repeated three times in preparation for DNA extraction. The extracted DNA was subsequently used in PCR reactions for further analyses of mutated genes in the patients. For the *DDX41* gene, the following primers were used: Forward - ATGGGTTAGGCCGGAAAAGGG, Reverse - TGACTCATCTGGGGGAGGAG. For the *PRPF8* gene, the primers used were Forward – AAGGAGACAATCCCCCGA and Reverse - TATAGGCCAGGTCAATGGCG.

### Flow cytometric analysis of mouse in vitro erythroid differentiation

Flow cytometric analysis of differentiation and enucleation of cultured mouse erythroblasts was conducted following previously established methods^19^. In brief, surface antigen labeling was performed using FITC or PE conjugated antibodies against CD71 and Ter119, along with their respective isotype controls. The stained cells were analyzed using a FACSCanto flow cytometer from Becton Dickinson. Additionally, propidium iodide was included to exclude non-viable cells from the analysis.

### G-quadruplexes immunofluorescence staining

Cells were washed with ice-cold serum-free Iscove’s Modified Dulbecco’s Medium (IMDM) and plated on poly-L-lysine-coated coverslips (BD Biosciences), followed by a 5-minute incubation at 37°C in a humidified incubator. After attachment, cells on the coverslip were washed with ice-cold PBS (pH 7.2), fixed in 4% paraformaldehyde for 15 minutes, and permeabilized with 0.1% Triton-X 100 in PBS for 10 minutes at room temperature. Following three PBS washes, cells were blocked with 3% BSA in PBS containing 0.05% Triton-X 100 for 1 hour at room temperature. Subsequently, the cells were stained with BG4 antibody and relevant secondary antibody for 1 hour each, followed by three cycles of PBS washing for 5 minutes each. Finally, the cells were mounted on glass slides using ProLong™ Diamond Antifade Mountant with DAPI (Invitrogen).

### Human CD34+ cell culture and differentiation

Human CD34+ cells (STEMCELL Technologies) were cultured in IMDM supplemented with 3% serum, 2% plasma, 10 µg/ml insulin, 3 IU/ml heparin, 200 µg/ml transferrin, 3 U/ml Epo, 10 ng/ml stem cell factor (SCF), 1 ng/ml interleukin-3 (IL-3), and 1% penicillin/streptomycin. The differentiation protocol involved a sequential withdrawal of cytokines. Initially, from day 0 to day 7, the culture included IL-3, EPO, and SCF, with medium exchanges every 2 days to maintain a cell concentration of approximately 1 × 10^5^ cells/ml. On day 7, a complete medium change was performed to eliminate IL-3. On day 10, SCF was withdrawn after a complete medium change. The cells were then maintained at 1 × 10^6^ cells/ml through regular medium changes until day 14. Starting from day 15, Epo was withdrawn through a complete medium change, and the cells were maintained at a concentration of 5 × 10^6^ cells/ml with regular medium changes until day 20. Cells were harvested at specific time points for analysis.

### CRISPR/Cas9 mediated DDX41 knockout in human CD34+ cells

The sgRNAs targeting DDX41 or scrambled sgRNA were cloned into the lentiviral vector lentiCRISPR v2 (Addgene, #52961, encoding Cas9) using the previously reported protocol^58^. The lentiviruses were produced in HEK293T cells following the manufacturer’s protocol (Invitrogen, MA, USA). For viral infection, roughly 3 × 10^7^ control or DDX41 sgRNA lentiviral particles were used to infect 5 × 10^5^ CD34+ cells. 14 hours after infection, cells were washed with PBS and cultured in a fresh medium. Cells were cultured for an additional 6 hours before follow-up studies.

### Cleavage Under Targets and Release Using Nuclease (CUT&RUN) assay

The following populations were subjected to CUT&RUN assays. HSPCs were isolated utilizing a negative selection methodology via the BD Biotin Mouse Lineage Panel kit (BD #559971). For the erythroid cells, biotin-conjugated Ter119+ antibody and magnetic streptavidin beads were used. The CUT&RUN assay was performed using the Cell Signaling Technology CUT&RUN Assay Kit (CST #86652). The harvested cells were incubated in a digitonin-containing buffer with primary antibodies at 4°C for 2 hours. pAG-MNase was then added and incubated at 4°C for 1 hour. To start the DNA digestion, pAG-MNase was activated with calcium chloride for 30 minutes at 4°C. The reaction was stopped using a stopping buffer. Under digitonin permeabilization, the digested DNA diffused into the supernatants. Finally, the digested DNA in the supernatants was purified using the spin column method and prepared for sequencing. Sequencing was performed at the Northwestern University NUseq core facility.

The initial data processing and differential peaks analysis were conducted by the NUseq core facility. Subsequently, the data analysis was performed utilizing the high-performance computer cluster at Northwestern Quest. Deeptools computeMatrix and plotHeatMap tools were employed to analyze and generate heat plots. The visualization of BigWig files was accomplished using the UCSC genome browser. Homer was used for motif enrichment analysis. Colocalization analysis was carried out by counting the read numbers within peak regions. Specifically, peak regions exhibiting more than 5 reads on DDX41 and G4 sites were classified as co-localized, while those not meeting this criterion were considered uniquely localized. Python was used to analyze and visualize the distribution of colocalization regions and peak regions. The sequencing data are available from the Gene Expression Omnibus database under accession code GSE254143.

### G4 pull-down assay

The oligonucleotides designed for G4 pull-down assay included sequences capable of forming G4 structures (G4-1: GATACTGCGAGTAGATACATGAAGGGAGGGCGCTGGGAGGAGGGATCT (derived from Kit1), G4-2: GATACTGCGAGTAGATACATGAACGGGCGGGCGCGAGGGAGGGGATCT (derived from Kit2), G4-3: GATACTGCGAGTAGATACATGAAGGCGAGGAGGGGCGTGGCCGGCATCT (derived from Spb1) and non-G4 controls (NC-1: GATACTGCGAGTAGATACATGAAGCAAGCACGCTGCAAGGAGCAATCT, NC-2: GATACTGCGAGTAGATACATGAACGCACGCACGCGAGCAAGCAGATCT, NC-3: GATACTGCGAGTAGATACATGAAGGCGACAAGGCACGTGGCCCACATCT), along with a complementary ssDNA sequence ATGTATCTACTCGCAGTATCATAC tagged with a 3’ biotin (BiosG). Oligonucleotides were initially dissolved to 100 μM in a buffer containing 10 mM Tris-HCl (pH 8.0) and 1 mM EDTA (TE buffer). For experimental use, they were further diluted to a 10 μM concentration. The ssDNA was mixed with either G4 or control oligonucleotides in a solution comprising 20 mM HEPES (pH 7.5), 250 mM KCl, and 1 mM DTT. This mixture was then heated at 95°C for 10 minutes, followed by a gradual cooling to 25°C at a rate of 0.1°C per second using a PCR machine.

For binding assays, high-affinity streptavidin magnetic beads (Pierce, Cat. No. 88817) were prepared in a binding buffer (10 mM Tris-HCl pH 8, 100 mM KCl, 0.1 mM EDTA, 1 mM DTT, 0.05% Tween-20) and incubated with the folded G4 oligonucleotides (0.5 nM oligonucleotides in 10 μl of streptavidin magnetic beads) for 2 hours at 4°C. Following two washes in the binding buffer to remove unbound oligonucleotides, the beads were incubated with 5 μg of DDX41 proteins for 1 hour at 4°C. Subsequent washes with binding buffers containing increasing concentrations of KCl (200–500 mM) were performed to remove non-specifically bound proteins. Proteins that remained bound to the beads were eluted by incubating for 5 minutes at 95°C in 2x SDS-PAGE loading buffer. The eluted proteins were then analyzed by SDS-PAGE and Western blotting to assess the binding interactions.

### G4 Fluorescence Resonance Energy Transfer (FRET) assay

G4 oligonucleotides were designed following previously published G4 oligo sequences^59,60^. Three oligonucleotides derived from the human *MYC* gene were generated. Each was conjugated with a FAM signal at the 5’-end and a BHQ group at the 3’-end. The sequences of these oligonucleotides are as follows: G4-1: 5’-FAM-TGGGGAGGGTGGGGAGGGTGGGGAAGGT-3’-BHQ, G4-2: 5’-FAM-GGGTGGGTTGGGTGGGG-3’-BHQ, and G4-3: 5’-FAM-GGGAGGGTTGGGTGGGG-3’-BHQ. To form G4 structures, these oligonucleotides were re-suspended in water and annealed in a tris-acetate solution containing 100 mM K+, with a total oligonucleotide concentration of 10 µM. The annealing was achieved by slow cooling from 95°C to room temperature. The resulting annealed G4 substrates were aliquoted into small tubes and stored at -20°C. Unfolding assays were conducted using a buffer composed of 20 nM oligonucleotides, 50 mM tris-acetate, 2.5 mM MgCl2, 0.5 mM DTT, and 50 mM KCl, along with varying concentrations of recombinant DDX41 protein. Fluorescence changes were monitored using an excitation wavelength of 490 nm and emission at 520 nm. The percentage of unfolding was calculated as: Unfolded G4 Percentage = (Sample G4 FAM Signal – Folded G4 FAM Signal) / (Unfolded G4 FAM Signal – Folded G4 FAM Signal)

### DDX41 protein expression and purification

Halo-tagged human DDX41 protein was expressed using the pJFT7-nHalo-DC vector containing the human DDX41 cDNA sequence (DNASU HsCD00947316). DH5α bacteria carrying the DDX41 plasmid were resuspended in 2 ml of LB medium supplemented with 50 µg/mL Ampicillin and 0.3% glucose. The suspension was incubated at 37°C overnight with vigorous shaking. Subsequently, the bacterial culture was diluted into 200 mL of LB medium containing 0.05% glucose and 0.05% rhamnose and incubated at room temperature for 1 day. The cells were then collected for protein purification using Promega HaloTag Protein Purification System. Briefly, the collected cells were first resuspended and lysed using lysozyme and sonicated in the presence of protease inhibitors to release the target protein. The targeted proteins were then bound to the resin. Subsequently, the bound protein was cleaved using TEV protease to release DDX41, while the Halo tag was removed. The purified proteins were then stored at -80°C for future use.

For the generation of DDX41 mutants (R525H and G530D), the DDX41 expression vector was modified using the GeneArt Site-Directed Mutagenesis System to introduce single-nucleotide mutations into the vector. The mutagenesis was carried out using the following primers: G530D Forward: GCTCGGGAAACACAGACATCGCCACTACCTT, Reverse: AAGGTAGTGGCGATGTCTGTGTTTCCCGAGC, R525H Forward: TTGGCCGCACCGGGCACTCGGGAAACACAGG, Reverse: CCTGTGTTTCCCGAGTGCCCGGTGCGGCCAA. Protein expression and purification of the mutant DDX41 variants followed the same method described for the wild-type DDX41.

### Quantitative RT-PCR of mouse 47s and 45s ribosome RNA

Total RNA was extracted from lysed cells using the Qiagen RNA extraction kit. Equivalent amounts of RNAs were employed for reverse transcription reactions with Takara PrimeScript RT Master Mix to generate cDNAs. The cDNAs were subsequently utilized as input for PCR reactions, performed using Sybr Green PCR mix. For the amplification of 45s rRNA, the following primer pairs were used: rn45sF1 (GCTGCGTGTCAGACGTTTTT) with rn45sR1 (AGAAAAGAGCGGAGGTTCGG), rn45sF2 (AGAGAACCTTCCTGTTGCCG) with rn45sR2 (AACTTTCTCACTGAGGGCGG), rn45sF3 (ATCGACACTTCGAACGCACT) with rn45sR3 (CACACGTCTGAACTTCGGGA), and rn45sF4 (TCCTTGTGGATGTGTGAGGC) with rn45sR4 (GGGAACATGGTCAAGCGAGA). For 47s rRNA amplification, the primer pairs included rn47sF1 (GGTGTCCAAGTGTTCATG) with rn47sR1 (CAAGCGAGATAGGAATGTCTTAC), rn47sF2 (AGAGAACCTTCCTGTTGCCG) with rn47sR2 (AACTTTCTCACTGAGGGCGG), and rn47sF3 (TCCTTGTGGATGTGTGAGGC) with rn47sR3 (GGGAACATGGTCAAGCGAGA).

### Chronic in vivo pyridostatin treatment in mice

To ensure prolonged exposure to pyridostatin (PDS), an ALZET osmotic pump (model 2006) was used. The pump allows for continuous drug delivery for up to 6 weeks. PDS was obtained from Millipore Sigma (Catalog No. SML2690), dissolved in saline, and loaded into the pumps. A final drug dosage of 3.5 mg/kg/day was achieved through this approach. As a control, saline was also loaded into separate pumps. These pumps were then subcutaneously implanted in wild-type mice. Following an 8-week period, we conducted analyses on the peripheral blood and bone marrow of the mice to evaluate the effects of chronic G4 accumulation stress.

### Single cell RNA sequencing

Mouse femoral bone marrow was extracted and immediately submitted for single-cell RNA sequencing (scRNA-seq) analysis. Cell viability assessments were conducted to ensure a minimum of 80% viability. The extracted samples were sent to the NUseq facility, where scRNA-seq experiments were conducted. Following scRNA-seq, the resultant raw sequence data in FASTQ format were processed using Cellranger on Northwestern University Quest High-Performance Computing Cluster. Subsequently, the processed scRNA-seq data were visualized and analyzed using the Cloupe browser. The sequencing data are available from the Gene Expression Omnibus database under accession code GSE254144.

### Induced pluripotent stem cell (iPSC)-derived human bone marrow organoid

The human bone marrow organoid differentiation was previously described^61^. In brief, iPSCs were purchased from StemCell Technologies (SCTi003-A). During differentiation, iPSCs were dissociated using EDTA when colonies reached approximately 100 μm in diameter. The iPSC aggregates were incubated overnight in mTeSR Plus medium (StemCell Technologies) enhanced with RevitaCell in 6-well Costar Ultra-Low Attachment plates (Corning, Cat#3471). Following this incubation, cells were gathered by gravitation in a 15 mL Falcon tube (Fisher Scientific, Cat#11507411) and resuspended in Phase I medium. This medium consisted of APEL2 (StemCell Technologies, Cat#05275) enriched with Bone Morphogenic Protein-4 (BMP4, Thermo Fisher Scientific, Cat#PHC9531), Fibroblast Growth Factor-2 (FGF2, StemCell Technologies, Cat#78134.1), and Vascular Endothelial Growth Factor-A (VEGF-165, StemCell Technologies, Cat#78159.1) at 50 ng/mL. The cells were then plated in 6-well Ultra-Low Attachment plates and incubated at 5% O2 for three days (days 0-3).

After this period, cell aggregates were collected by gravitation and suspended in Phase II medium for an additional 48 hours (days 3-5). The Phase II medium included APEL2 supplemented with BMP-4, FGF2, and VEGFA at 50 ng/mL, along with human Stem Cell Factor (hSCF, StemCell Technologies, Cat#78062) and Fms-like tyrosine kinase-3 Ligand (Flt3, StemCell Technologies, Cat#78009) at 25 ng/mL. On day 5, cells were again collected by gravitation for hydrogel embedding. The hydrogels, composed of 60% collagen and 40% Matrigel, were prepared on ice following the manufacturer’s instructions. This included Reduced Growth Factor Matrigel (Corning, Cat#354230) supplemented with 1 mg/mL human collagen type I (Advanced Biomatrix, Cat#5007) and type IV (Advanced Biomatrix, Cat#5022). A 0.5 mL cell-free base layer was first laid and allowed to polymerize for 2 hours, followed by another 0.5 mL layer containing the cell aggregates, also left to polymerize for 2 hours at 37°C and 5% CO_2_. The fully polymerized gels with cell aggregates were then supplemented with Phase III media. This media included VEGFA at either 50ng or 25 ng/mL, VEGFC (where applicable) at 50 or 25 ng/mL, FGF2, BMP4, hSCF, Flt3, Erythropoietin (EPO, StemCell Technologies, Cat#78007), Thrombopoietin (TPO, StemCell Technologies, Cat#78210), Granulocytic Colony-Stimulating Factor (G-CSF, StemCell Technologies, Cat#78012), at 25 ng/mL, and Interleukin-3 (IL3, StemCell Technologies, Cat#78194) and Interleukin-6 (IL6, StemCell Technologies, Cat#78050) at 10 ng/mL. The media was refreshed every 72 hours. On day 12, the organoids were scooped from hydrogels and transferred to 96-well ultralow attachment plates (Thermo Fisher, Cat#174925) for further culturing. The media was refreshed at 1:1 ratio every 72 hours. Organoids are fully matured on day 21 for downstream experiments.

Confocal imaging of the whole-mount organoids was conducted on a Nikon AXR confocal system, utilizing a 20X water immersion objective (CFI Apo LWD Lambda S 20XC WI). Organoids were fixed in 4% PFA and subsequently immunostained with primary and secondary antibodies: anti-human CD45, 1:500 (eBioscience, 14-9457-82); anti-CD140b, 1:200 (Abcam, ab215978),anti-CD71, 1:250 (eBioscience, 14-0719-82); anti-CD235a, 1:200 (PA5-141179); biotin-UEA1, 1:200 (Vector Lab, B0-1065-2), AF647 conjugated goat anti-rabbit IgG (H+L) cross-adsorbed secondary antibody; AF568 conjugated streptavidin; AF488 conjugated goat anti-mouse IgG (H+L) cross-adsorbed secondary antibody. All secondary antibodies were used at 1:300 dilution. Following immunostainings, organoids were dehydrated in ethanol of different concentrations (50%, 70%, 90%, 100%) before tissue clearance with Ethyl Cinnamate and subsequent imaging. Z-stack confocal images were rendered with Imaris 10.0 (Oxford Instruments).

### Engraftment of CD34+ HSPCs

CD34+ hematopoietic stem and progenitor cells (HSPCs) were purchased from StemCell Technologies (Cat#70002). For engraftment, CRISPR/Cas9-edited CD34+ cells were stained with CellVue Claret Far Red Membrane Label (Sigma-Aldrich, Cat#MINCLARET-1KT) following the manufacturer’s instructions. 5×10^4^ CellVue-labeled CD34+ cells were suspended in 200 µL StemPro-34 medium (Gibco, Cat#10639011) and combined with 100 µL Matrigel (Corning, Cat#CLS356237) on ice. Organoids on day 18 post-differentiation were deprived of their cell culture medium and exposed to 30 µL of this cell suspension in ultra-low attachment 96-well plates (Thermo Fisher, Cat#174927), followed by a 30-minute incubation at 37°C. Afterward, 150 µL of StemPro-34 medium containing SCF, FLT3, TPO, EPO, and IL3 (10 ng/mL each) was added to support engraftment. The engrafted organoids were evaluated via flow cytometry or imaging 3 days after engraftment. We pooled 10 engrafted organoids for each flow cytometry analysis to mitigate the variability between individual organoids.

## Statistics

Results are expressed as mean ± SEM unless otherwise indicated. Statistical comparisons between two groups were performed with two-tailed unpaired Student’s t tests, and the comparison among multiple groups was evaluated with 1-way ANOVA tests using GraphPad Prism version 9.0 software.

## Acknowledgments

We thank Ching Man Wai and Matthew Schipma from the NUSeq Core for their help with the sequencing studies. We thank Susan Ross for providing Ddx41 floxed mice and Nicolae Valentin David for HBBCre mice. This work was supported by the National Institute of Diabetes and Digestive and Kidney Disease (NIDDK) grant R01-DK124220 (P.J.), National Heart, Lung, and Blood Institute (NHLBI) grant R01-HL148012 (P.J.), R01-HL150729 (P.J.), R01-HL169507 (P.J.)., R35-HL171168 (L.B.), National Cancer Institute (NCI) grant R00-CA248835 (V.S.). H.B. is a recipient of the F32 Ruth L. Kirschstein Postdoctoral Individual National Research Service Award (F32-HL170648). K.R. is a recipient of the Leukemia & Lymphoma Society Career Development Program Award. P.J. is a scholar of the Leukemia & Lymphoma Society.

## Author contributions

H.B., K.R., P.W., E.L., X.H., J.Y., I.A., K.T., and Y.T. performed the experiments and interpreted data. H.B., E.T.B, and W.W. performed the CUT&RUN data analyses. H.B., K.R., L.G., Y.L., V.S., L.B., M.S., and P.J. analyzed the data. H.B. and P.J. designed the experiments, interpreted data, and wrote the manuscript.

## Supplemental figures and figure legends

**Supplemental Figure 1.**
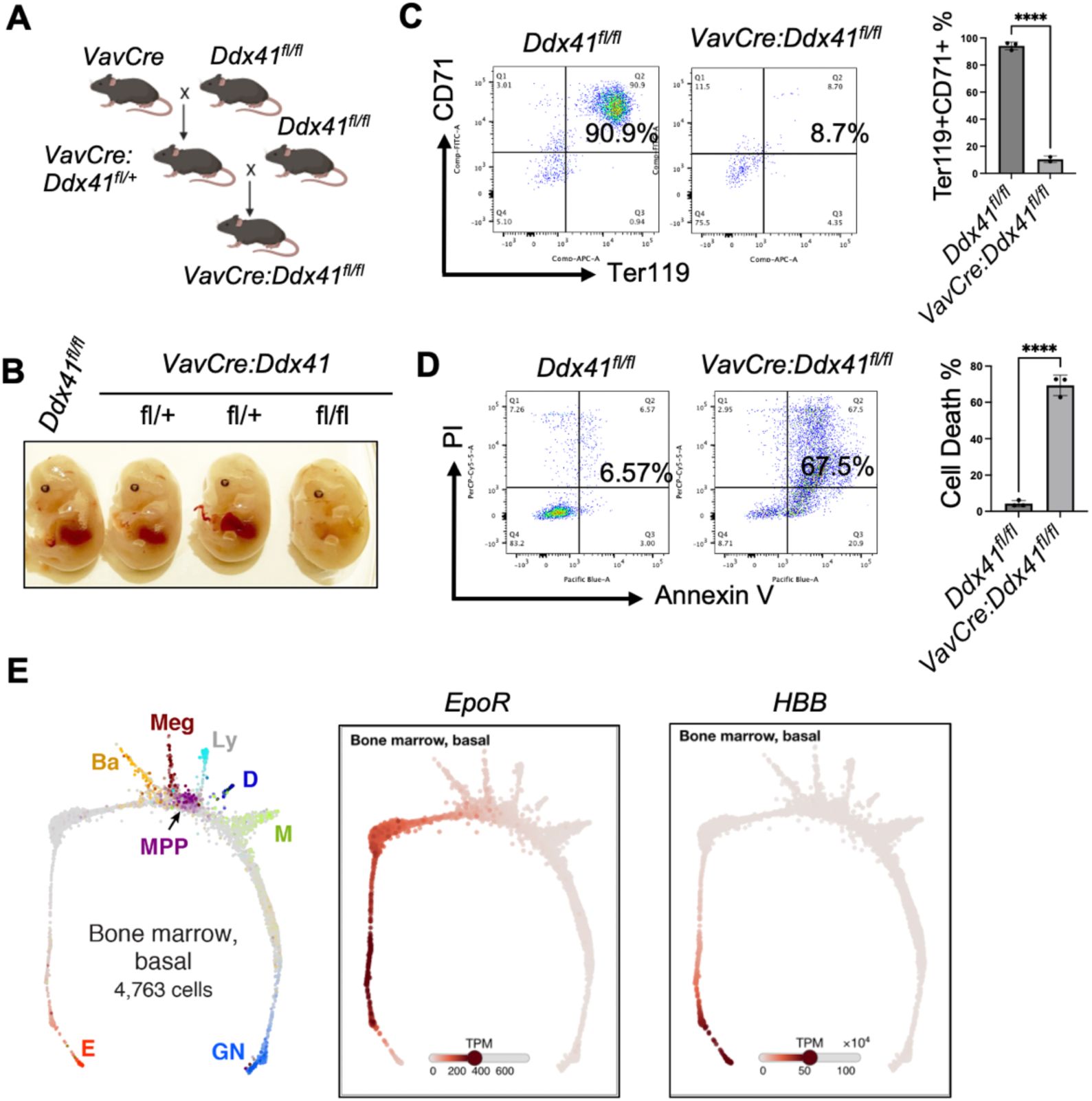
Ddx41 is essential for hematopoiesis. (A) Schematic representation of the breeding strategy to generate hematopoietic-specific *Ddx41* knockout mice. (B) Representative pictures of E14.5 embryos with the indicated genotypes. (C-D) Ter119 negative fetal liver cells from E14.5 embryos of the indicated mice were cultured in an Epo-containing medium for 48 hours. The differentiation status was evaluated by flow cytometry using CD71 and Ter119 markers. Cell viabilities were assessed using propidium iodide (PI) and Annexin V staining. Quantifications are on the right. (E) Expression patterns of EpoR and HBB during hematopoiesis. The left UMAP panel displays cells from various lineages and stages. The middle and right panels illustrate the expression pattern of EpoR and HBB, respectively. Figures were generated using the online tool accessible at https://kleintools.hms.harvard.edu/paper_websites/tusi_et_al/.

**Supplemental Figure 2.**
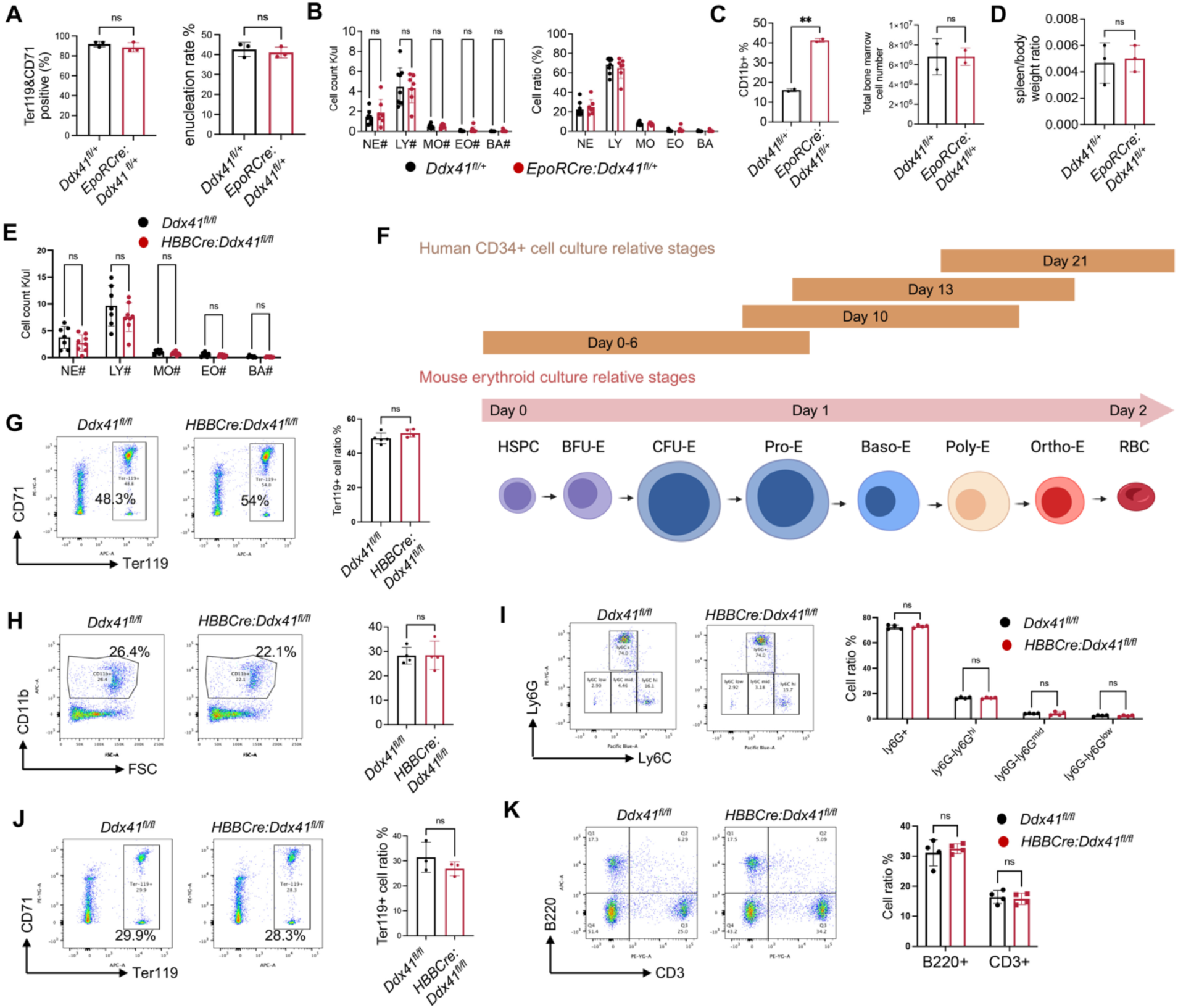
Ddx41 is differentially required at different stages of terminal erythropoiesis. (A) Ter119-negative cells from the E13.5 fetal liver were purified from the indicated mice and cultured in Epo medium for 2 days. Cell differentiation and enucleation were measured by flow cytometry. (B) Leukocyte count of indicated mice at 2 months old. (C) Flow cytometry assay of bone marrow myeloid cells from mice in B using CD11b as a marker. Quantification of B and total bone marrow cell number is on the right. (D) Spleen/body weight ratio of indicated mice in A. (E) Leukocyte count of indicated mice at 2 months old. (F) Schematic diagram of the corresponding stages of erythropoiesis in the mouse and human in vitro culture systems. (G) Flow cytometry assay of bone marrow erythroid cells from mice in E using CD71 and Ter119 as markers. Quantification is on the right. (H-K) Flow cytometry assays of bone marrow myeloid (H), bone marrow myelomonocytic (I), bone marrow erythroid (J), and spleen lymphoid (K) populations from mice in E. Quantifications are on the right. At least three independent samples were used in each experiment. P-values were determined with 2 tailed t tests. *p<0.05, **p<0.01, ***p<0.001, ****p<0.0001.

**Supplemental Figure 3.**
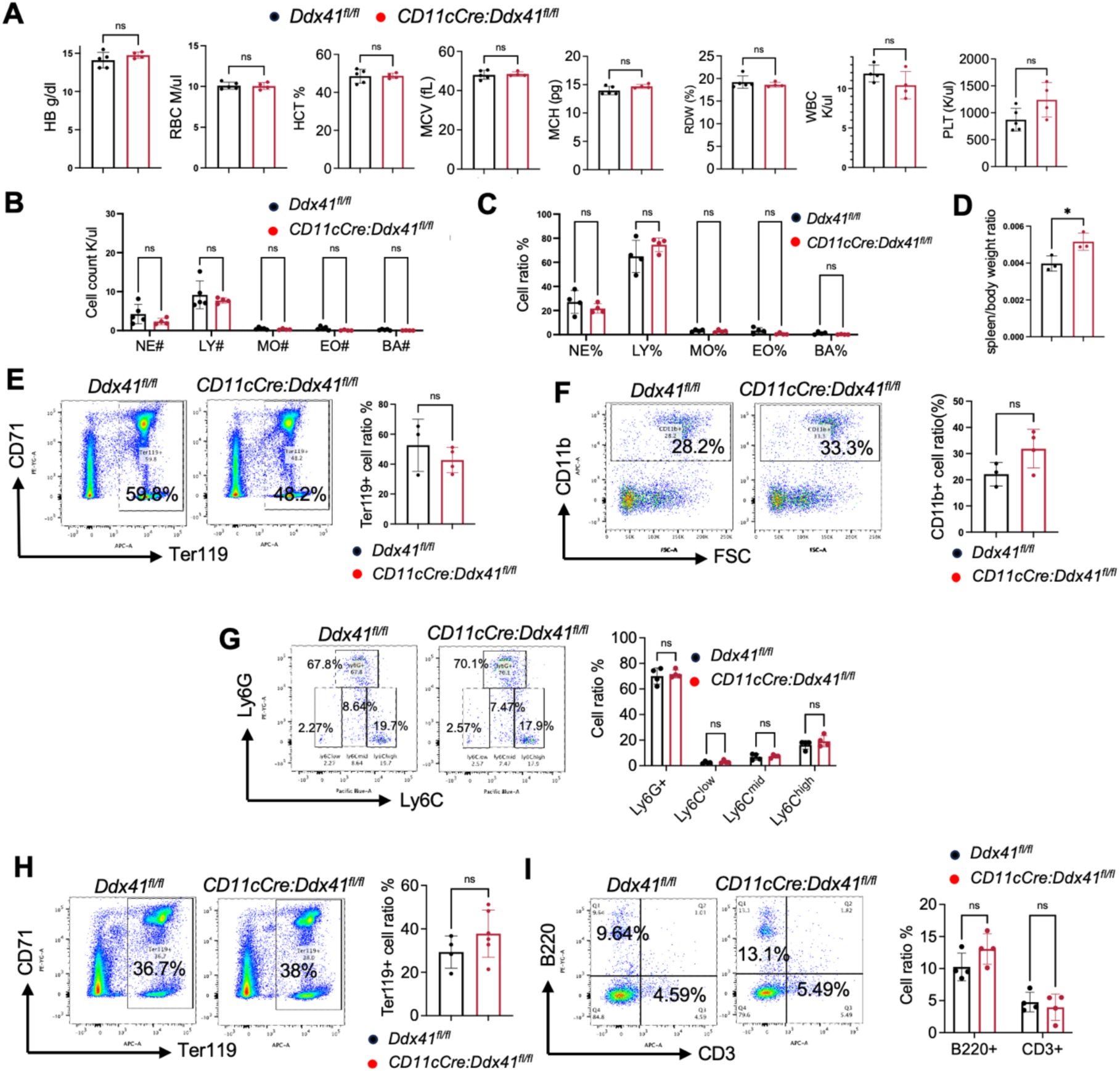
Ddx41 is dispensable for dendritic cell differentiation. (A) Complete blood count of indicated mice at 2 months old. (B-C) Leukocyte absolute number (B) and percentage (C) of mice from A. (D) Spleen/body weight ratio of indicated mice in A. (E-I) Flow cytometry assays of bone marrow erythroid (E), bone marrow myeloid (F), bone marrow myelomonocytic (G), spleen erythroid (H), and spleen lymphoid (I) populations from mice in A. Quantifications are on the right. Each panel represents data obtained from a minimum of three independent samples. P-values were determined with 2 tailed t tests. *p<0.05, **p<0.01, ***p<0.001, ****p<0.0001.

**Supplemental Figure 4.**
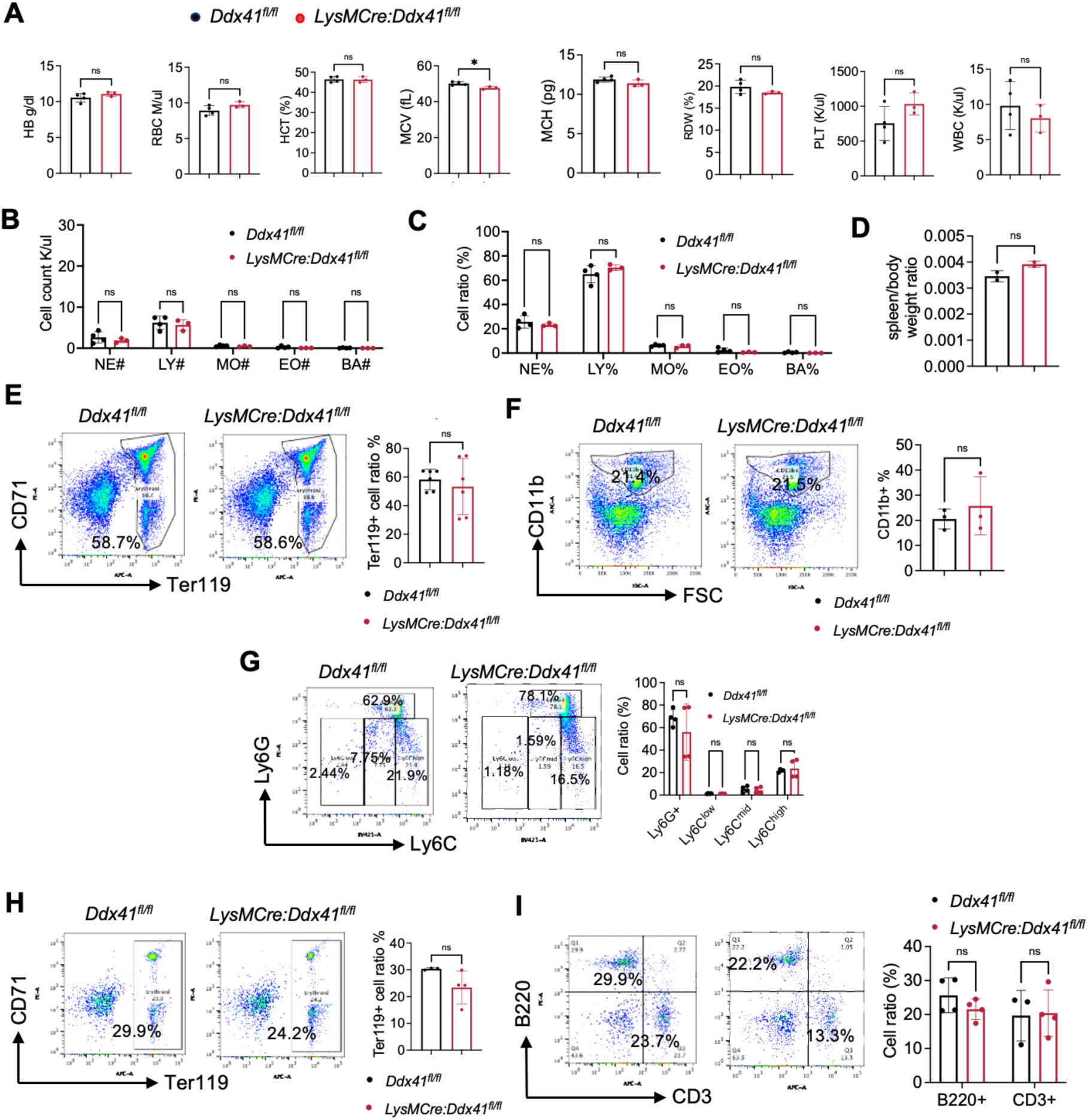
Ddx41 is dispensable for monocytic cell differentiation. (A) Complete blood count of indicated mice at 2 months old. (B-C) Leukocyte absolute number (B) and percentage (C) of mice from A. (D) Spleen/body weight ratio of indicated mice in A. (E-I) Flow cytometry assays of bone marrow erythroid (E), bone marrow myeloid (F), bone marrow myelomonocytic (G), spleen erythroid (H), and spleen lymphoid (I) populations from mice in A. Quantifications are on the right. Each panel represents data obtained from a minimum of three independent samples. P-values were determined with 2 tailed t tests. *p<0.05, **p<0.01, ***p<0.001, ****p<0.0001.

**Supplemental Figure 5.**
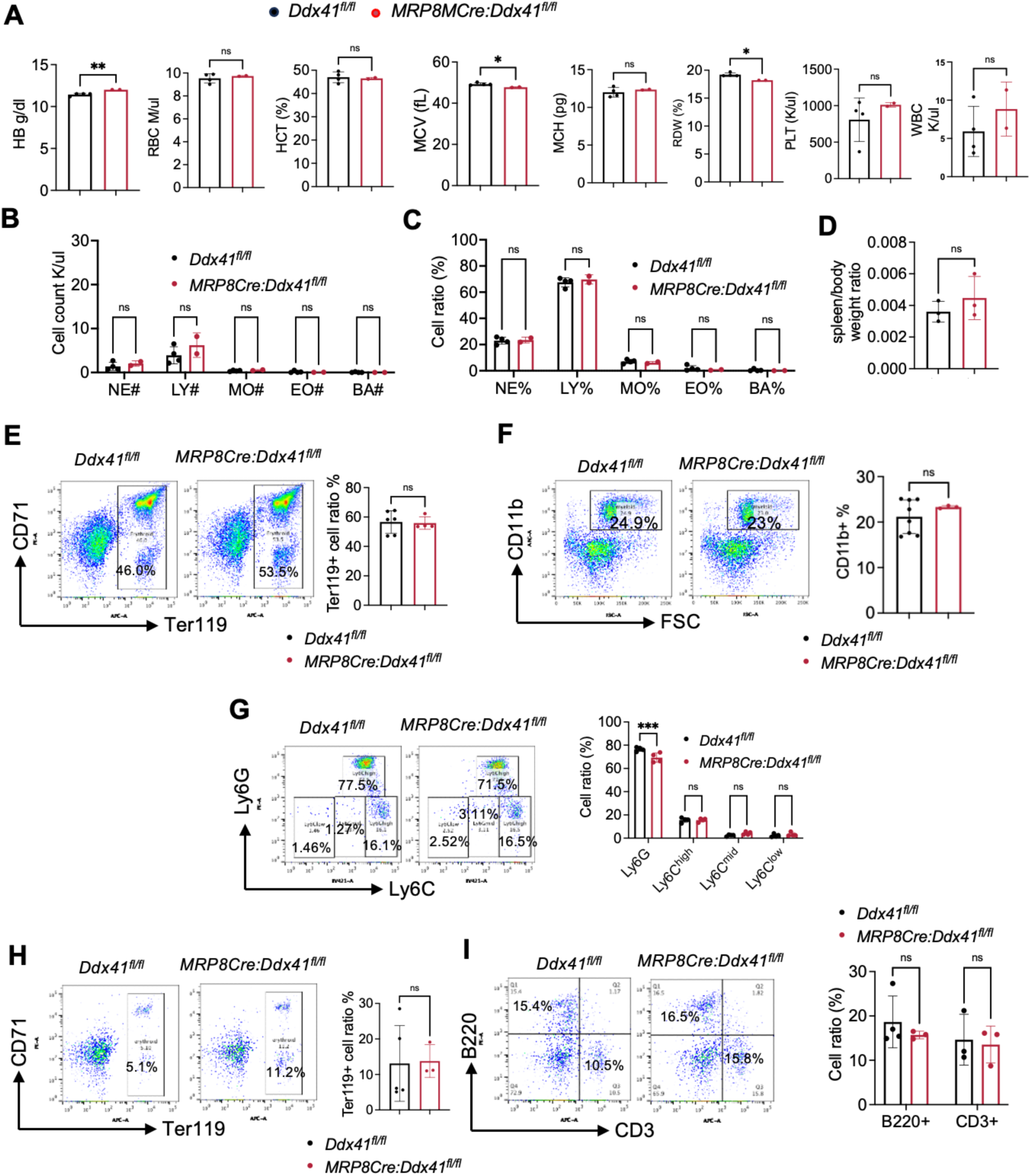
Ddx41 is dispensable for myeloid cell differentiation. (A) Complete blood count of indicated mice at 2 months old. (B-C) Leukocyte absolute number (B) and percentage (C) of mice from A. (D) Spleen/body weight ratio of indicated mice in A. (E-I) Flow cytometry assays of bone marrow erythroid (E), bone marrow myeloid (F), bone marrow myelomonocytic (G), spleen erythroid (H), and spleen lymphoid (I) populations from mice in A. Quantifications are on the right. Each panel represents data obtained from a minimum of three independent samples. P-values were determined with 2 tailed t tests. *p<0.05, **p<0.01, ***p<0.001, ****p<0.0001.

**Supplemental Figure 6.**
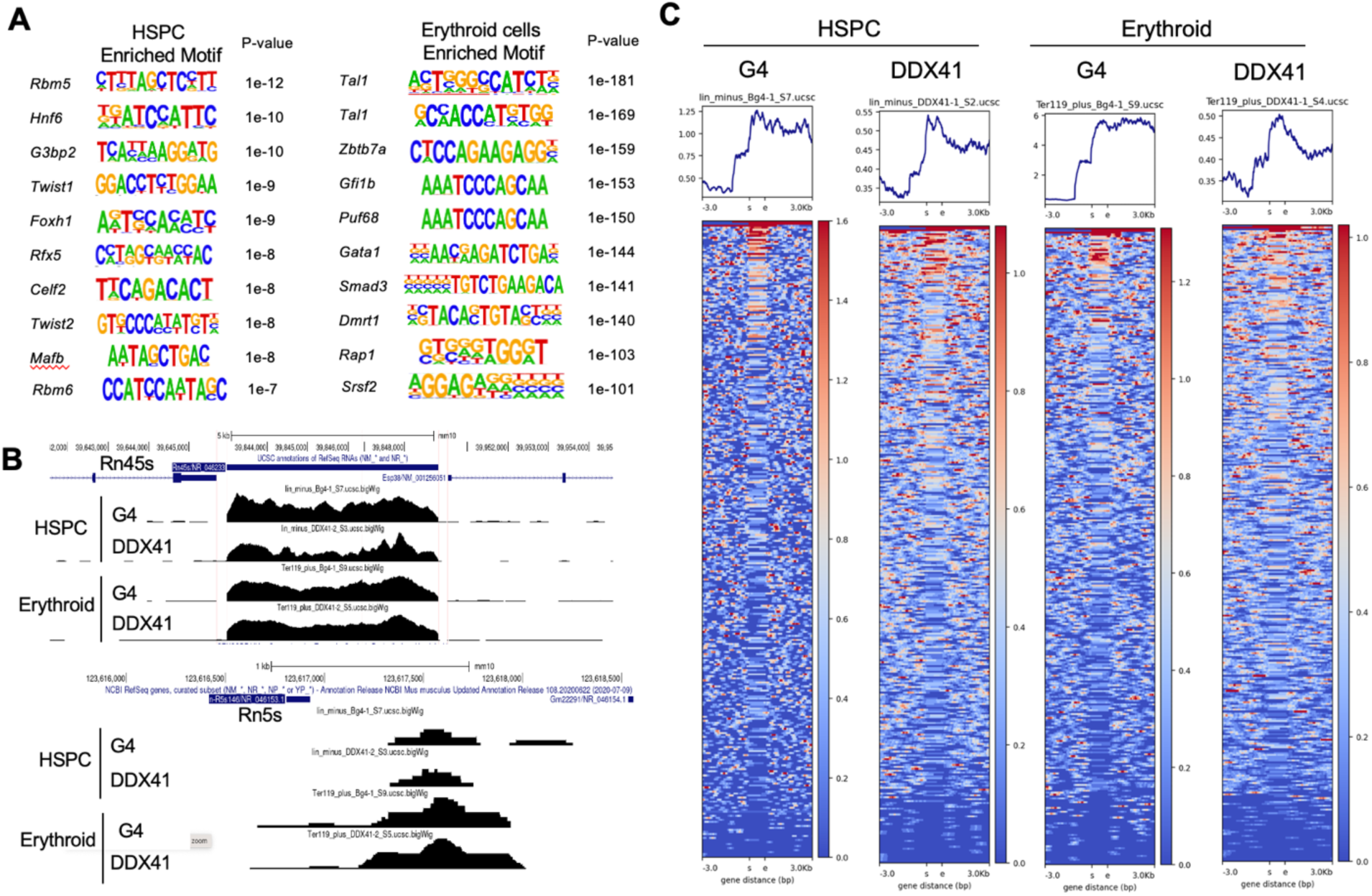
Ddx41 colocalizes with G4 at the erythroid genome level. (A) Motif enrichment analysis of colocalized G4 and Ddx41 sites in mouse HSPC and erythroid cells. Motif enrichment analysis was conducted using the Homer Motif analysis tool, with no GC weight applied. (B-C) Enrichment of G4 and Ddx41 binding sites in rDNAs in HSPC and erythroid cells. Panel B displays exemplary data with G4 and Ddx41 binding peaks enriched in Rn45s and Rn5s genes. Panel C presents a heat map illustrating the localization of G4 and Ddx41 within rDNAs. The rDNA sequences were retrieved from the UCSC genome browser using the selection criteria of matching gene type ’*rRNA’ from the mm10 (VM25) assembly.

**Supplemental Figure 7.**
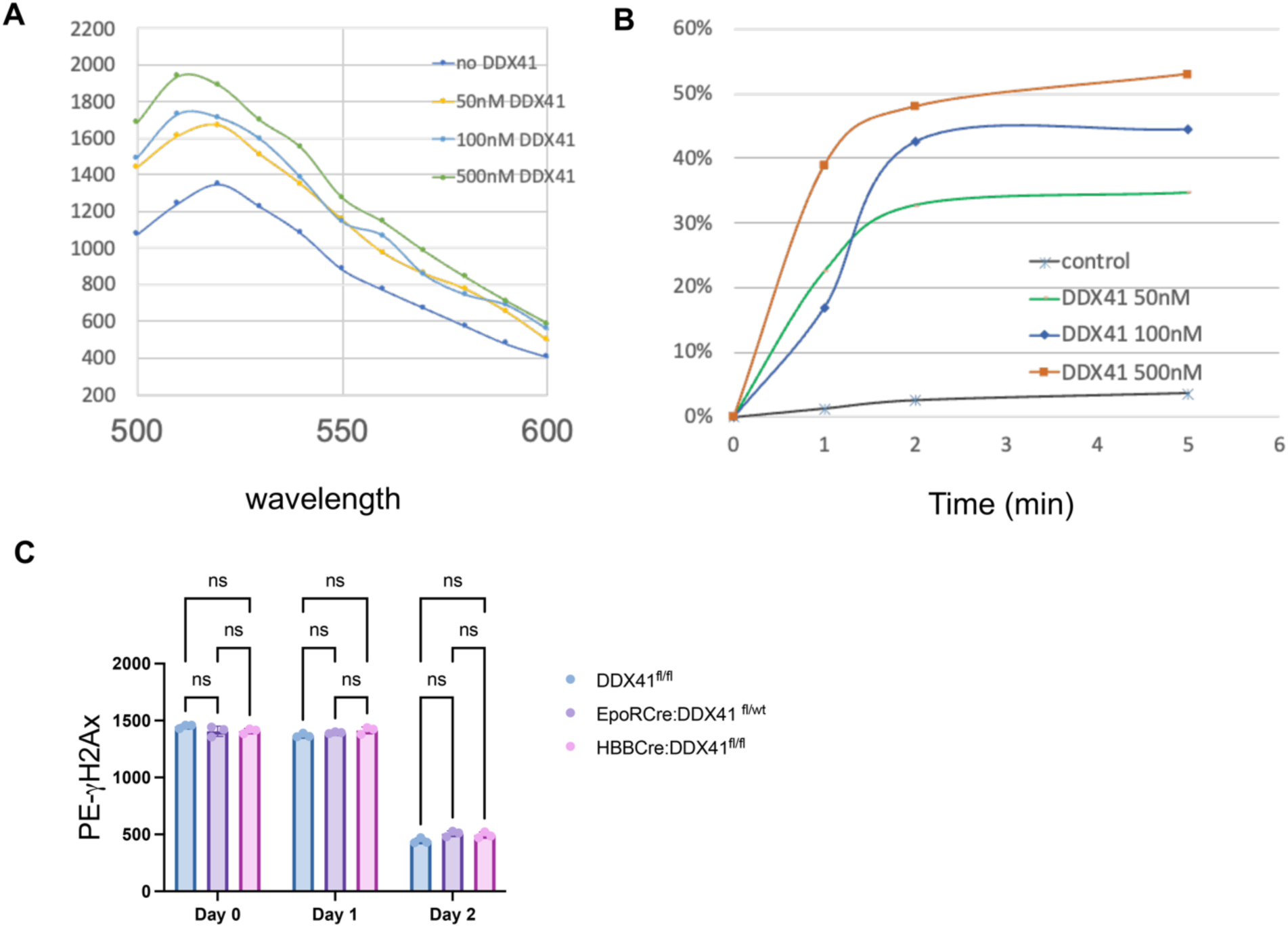
DDX41 dose-dependently dissolves G4. (A) Dose-dependent increase of FAM signal with increased concentration of recombinant human DDX41 protein. (B) Time course of DDX41-mediated G4 dissolving activity. (C) Lineage-negative cells from the bone marrow of the indicated mice were purified and cultured in Epo medium for 2 days. The levels of γ-H2AX were measured and quantified using flow cytometry.

**Supplemental Figure 8.**
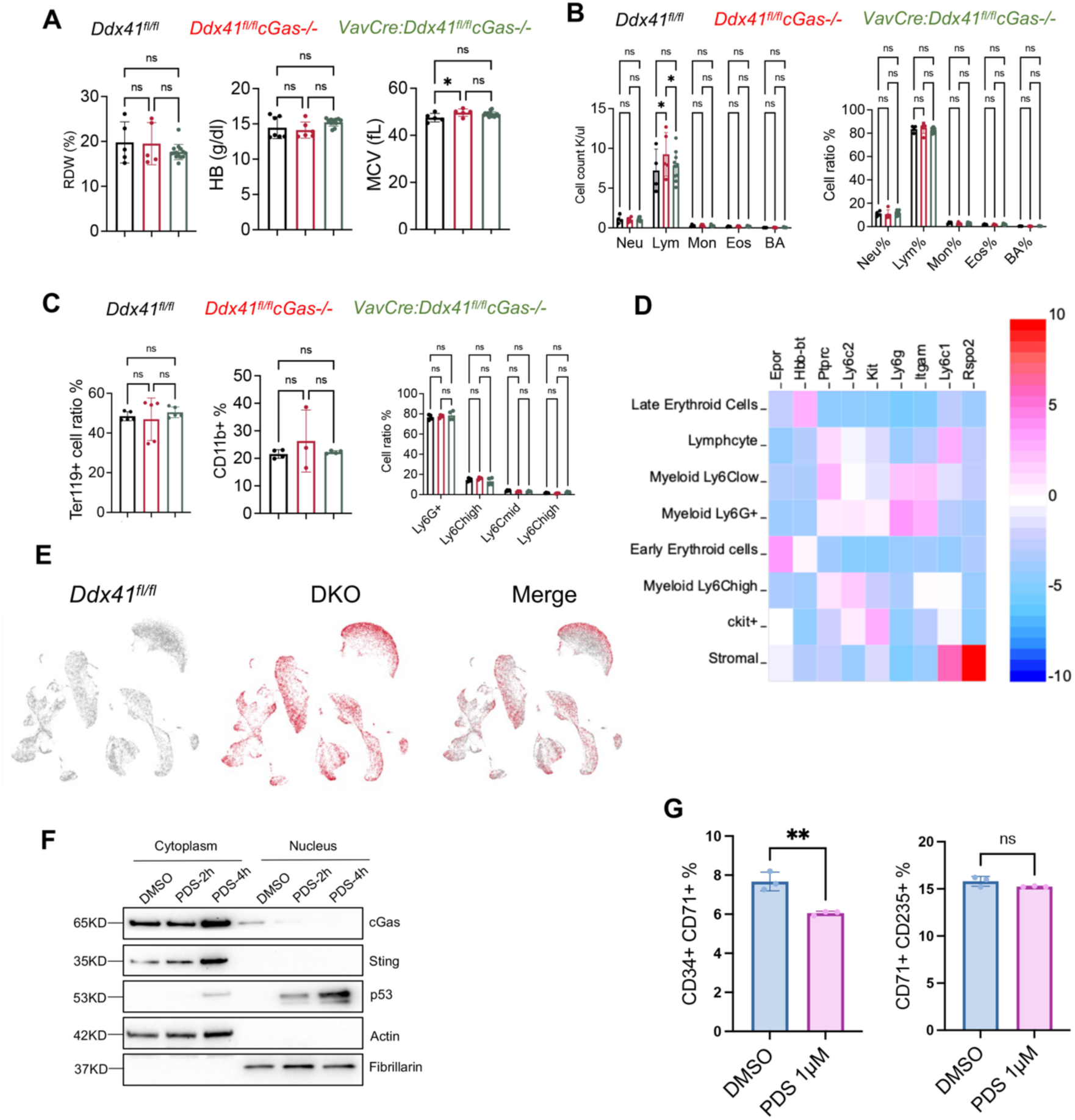
cGas deficiency rescues the embryonic lethality of hematopoietic specific Ddx41 knockout mice. (A) Red cell indices of the indicated mice at 2 months old. (B) Leukocyte absolute number (left) and percentage (right) of mice from A. (C) Quantification of flow cytometry assays of bone marrow erythroid (left), myeloid (middle), and myelomonocytic (right) from mice in A. (D) Specific marker genes in the identified cell clusters in the scRNA sequencing data. (E) Uniform Manifold Approximation and Projection (UMAP) plots showing the distribution and overlapping of different cell populations in the bone marrow of indicated mice from A. (F) Western blotting assays of indicated proteins in the cytoplasmic and nuclear fractions of bone marrow erythroid cells treated with DMSO or PDS (1 μM) for the indicated amount of time. Lineage-negative cells from the bone marrow of wild-type mice were cultured in Epo medium for 1 day, followed by the treatment. (G) Human bone marrow organoids were treated with PDS for 24 hours followed by flow cytometry assays of the indicated cell types. All the error bars represent the SEM of the mean. The comparison among multiple groups was evaluated with 1-way ANOVA tests. ns: non-significant. *p<0.05.

**Supplemental Table 1.**

| Lineage-specific Cre system | Expression lineage | Earliest knockout stage | Viable homozygous mutant? | Homozygous fetuses acquired or not? |
| --- | --- | --- | --- | --- |
| ***Vav-Cre*** | HSCs and HSPCs | HSCs (~E13.5) | No | Yes |
| ***EpoR-Cre*** | Erythrocytes | Primitive erythroblasts, early terminal erythroblasts | No | Yes |
| ***HBB-Cre*** | Erythrocytes | Late terminal erythroblasts | Yes | Yes |
| ***CD11c-Cre*** | Dendritic cells (DCs) | Pre-cDCs, pre-pDCs | Yes | Yes |
| ***MRP8-Cre*** | Granulocytes, granulocyte/macrophage progenitors (GMPs) | Myelocyte stage | Yes | Yes |
| ***LysM-Cre*** | Monocytes, mature macrophages, and granulocytes | Immature monocytes | Yes | Yes |

